# Differential higher-order superassembly of HECT-type UBE3 ligases controlled by calcium signals

**DOI:** 10.64898/2026.02.09.704917

**Authors:** Xin Luo, Miyun Shi, Jiao Hu, Deyao Yin, Shitao Zou, Qihao Wang, Xuemei Li, Guopeng Wang, Yanzhen Hou, Youdong Mao

## Abstract

The yeast E4 ligase Hul5 transiently associates with the proteasome during heat shock and proteotoxic stress to maintain cytosolic protein quality control. Two of its human orthologs, UBE3B and UBE3C, have specialized to safeguard mitochondria and inherit Hul5 function, respectively. We determined cryo-electron microscopy structures of full-length human UBE3B and UBE3C in complex with calmodulin at 2.9-3.5 Å in calcium-free and calcium-saturated conditions. Calmodulin acts as an inter-protomer molecular glue clamping UBE3B into ring-shaped anti-parallel homodimers or asymmetric trimers, and remodels UBE3C into a conformation competent for proteasome association. Calcium binding to calmodulin promotes disassembly of these higher-order complexes, toggling UBE3B/C from E4 to E3 activity. These findings reveal how calmodulin regulates higher-order architectures of E3/E4 ligases to rewire the ubiquitin-proteasome system via calcium signaling.

---

Eukaryotic survival depends on the robustness of cellular quality control networks under metabolic fluctuations and proteotoxic stress (*1*), with the ubiquitin–proteasome system (UPS) as a central component (*2*). E3 ubiquitin ligases function as decision-makers in protein quality control by targeting substrates for proteasomal degradation or selective autophagic clearance (*3, 4*). Many E3 ligases assemble into higher-order organization, typically homodimers, to orchestrate allosteric regulation of catalytic activities and support multivalent substrate recognition (*5, 6*). These E3 super-assemblies allow quality control systems to tune their degradative capacity in response to changing levels of stress (*7*).

The Homologous to the E6-AP Carboxyl Terminus (HECT)-type E3 ubiquitin ligases UBE3B and UBE3C, two mammalian orthologs of the yeast ligase Hul5, are key guardians of protein quality control under stress conditions (*8*). UBE3B localizes to the mitochondria–cytosol interface, where it safeguards mitochondrial integrity and suppresses ferritinophagy by ubiquitylating the autophagy cargo receptor NCOA4 (nuclear receptor coactivator 4) (*9, 10*). UBE3B also maintains cellular homeostasis by removing the aberrant accumulation of signaling effectors (*11–14*). By contrast, UBE3C and Hul5 associate with the proteasome to enhance proteasomal processivity by orchestrating their E4 ubiquitin-chain extension activity and clear degradation-resistant substrates (*8*). UBE3C further modulates autophagic activity by ubiquitylating autophagy regulators such as ATG4B (autophagy related 4B cysteine peptidase), thereby tuning autophagic flux (*15*). Reflecting these central functions, loss of UBE3B or UBE3C causes neurodegenerative or neurodevelopmental disorders, including Kaufman oculocerebrofacial syndrome (KOS) (*16*) and Distal Hereditary Motor Neuropathy (DHMN1) (*17*), and their dysregulation is linked to the progression of multiple cancers (*11, 12, 18–20*). Together, UBE3B and UBE3C play central roles in protein quality control and coordinating UPS with autophagy.

Intracellular calcium (Ca^2+^) functions as a second messenger that orchestrates essential physiological processes ranging from neuronal synaptic transmission to metabolic adaptation (*21*). Cells process Ca^2+^ signals through calmodulin (CaM), a primary calcium sensor that undergoes Ca^2+^-dependent conformational rearrangements to regulate downstream effectors (*22*). Emerging evidence indicates that Ca^2+^-dependent CaM binding can remodel intermolecular interfaces and drive transitions between monomeric and dimeric or oligomeric states, creating regulatory switches that modulate enzyme activity and substrate specificity (*23, 24*). Despite their distinct subcellular localizations and substrate specificity, UBE3B and UBE3C share a conserved N-terminal IQ motif that serves as a CaM recognition site. Previous biochemical studies on UBE3B showed that CaM binding to this IQ motif suppresses its E3 ligase activity, whereas elevated Ca^2+^ levels induce CaM dissociation and relieve this inhibition (*9*). However, how Ca^2+^-dependent CaM binding controls the conformational dynamics and higher-order superassembly of UBE3B and UBE3C remain unknown.

Here we combine high-resolution cryo-electron microscopy (cryo-EM), biochemical studies and AlphaFold3 (AF3) modelling (*25*) to define the structural basis of Ca^2+^/CaM-mediated regulation of UBE3B and UBE3C functions and their interactions with important substrates, including the oncogenic transcription factor MYC (*11*) and the oxygen-sensing protein HIF-2α (hypoxia-inducible factor-2α) (*12*). Our analyses reveal a striking architectural divergence in the higher-order superassemblies of UBE3B and UBE3C orchestrating distinct responses to Ca^2+^ signals. UBE3B forms CaM-mediated oligomers potentially amplifying Ca^2+^-dependent responses, whereas UBE3C instead utilizes a highly flexible N-terminal tail to assemble with the 26S proteasome. Our findings elucidate how a conserved Ca^2+^-sensing mechanism in the ubiquitin-proteasome system has been evolved to regulate E3/E4 ubiquitin ligase activities in essential cellular processes via diverse higher-order superassemblies.

## Structures of human UBE3B–CaM complexes

To obtain stable complexes suitable for structural analysis, we co-expressed full-length human UBE3B and UBE3C with CaM in Expi293F cells, respectively. Subsequent tandem affinity purification followed by size-exclusion chromatography (SEC) yielded homogeneous UBE3B–CaM and UBE3C–CaM samples. The SEC profiles and SDS-PAGE analysis of the peak fractions demonstrated that both UBE3B and UBE3C co-elute with CaM as stable complexes (fig. S1, A to D). We further verified that the purified complexes for cryo-EM studies were enzymatically active using an *in vitro* ubiquitylation assay (fig. S1E).

To explore the structural organization of UBE3B, we performed cryo-EM analysis of a native, crosslinking-free sample in 2 mM EGTA, which revealed co-existing and monomeric, dimeric and trimeric populations. Reconstructions of the full-length UBE3B–CaM monomer, dimer and trimer were obtained at average resolutions of 2.9, 3.3 and 4.0 Å, respectively. We overcame the preferred orientation problem for the dimer by combining tilted and untilted datasets (Fig. 1, B to E; fig. S2, A to D). The cryo-EM map of the trimer reveals an asymmetric superassembly architecture in which an additional protomer binds to a ring-shaped dimer (Fig. 1D). Consistent with our cryo-EM reconstructions, SDS-PAGE and mass photometry analysis of the purified UBE3B after crosslinking treatment revealed distinct high-molecular-weight bands, corresponding to dimers and trimers (fig. S1, F and G).

**Fig. 1.**
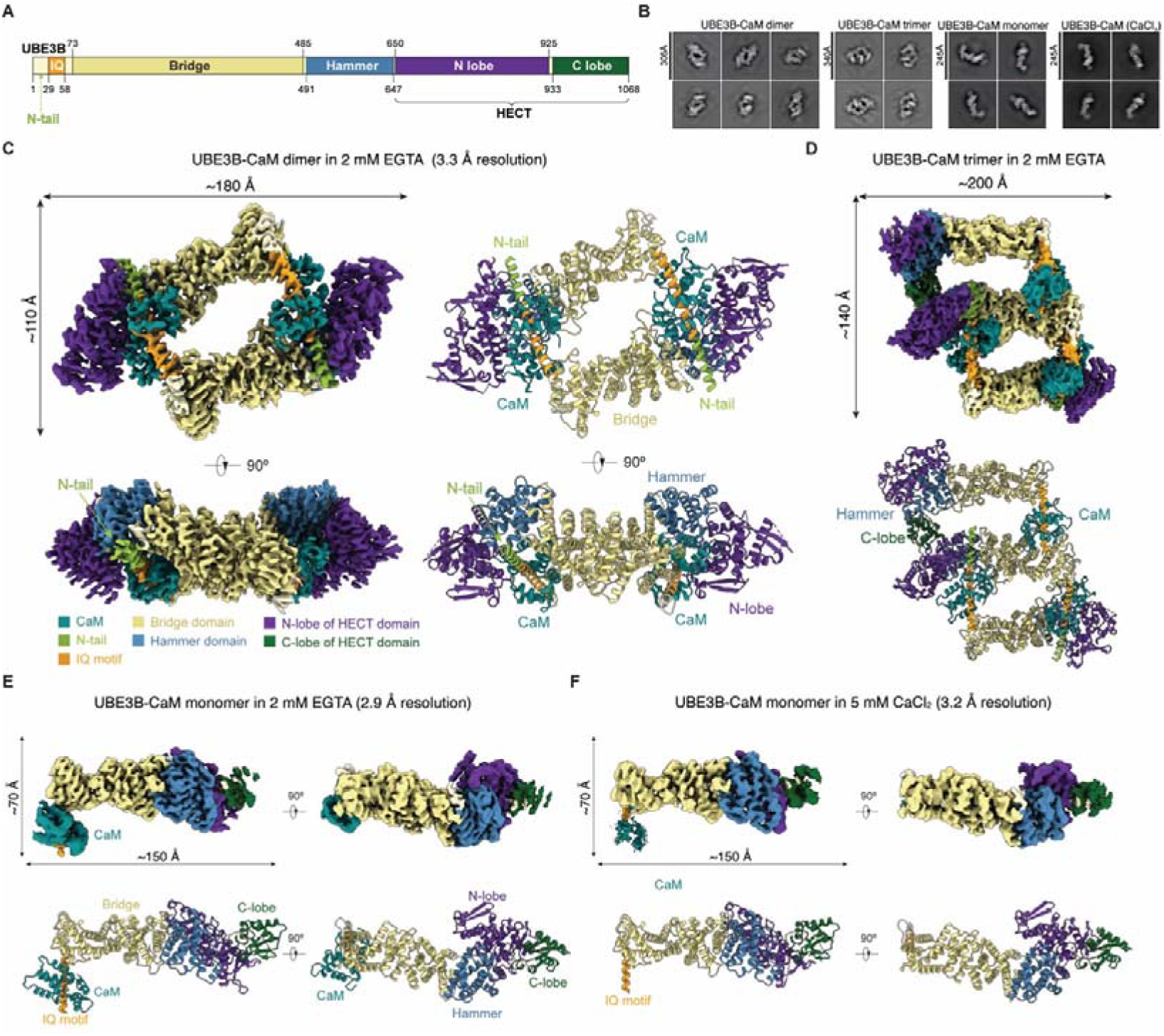
Structural basis for CaM-mediated high order assembly of UBE3B. (**A**) Domain organization of human UBE3B, including the IQ motif, N-tail, Bridge, Hammer, and HECT domain. (**B**) Representative 2D class averages of UBE3B–CaM oligomers and monomer in 2 mM EGTA (from left to middle right) and in 5 mM CaCl_2_ (monomer, right). Box sizes are labeled in Å. (**C**) Cryo-EM density map (left) and atomic model (right) of the UBE3B–CaM dimer (3.3 Å resolution) in the presence of 2 mM EGTA, shown from different orientations. Maps and models are color-coded by domain: HECT N-lobe (purple), Bridge (yellow), Hammer (blue), and bound CaM molecules (dark cyan). Note that the HECT C-lobe is not modeled due to conformational variability. (**D**) Cryo-EM density map (top) and atomic model (bottom) of UBE3B trimer in the presence of 2 mM EGTA with domains colored as in (C). (**E** and **F**) Cryo-EM density map (top) and atomic model (bottom) of UBE3B–CaM monomer in the presence of 2 mM EGTA at 2.9 Å resolution (E) and in 5 mM CaCl_2_ at 3.2 Å resolution (F). HECT C-lobe (dark green) rigid-body fitted from an AlphaFold 3 (AF3) model. The Ca^2+^-bound CaM is omitted in (F) due to lack of resolvable density. Domains are colored as in (C).

The domain organizations of UBE3B and UBE3C feature a conserved N-terminal IQ motif, which mediates the interaction with CaM (Figs. 1A and 2A). The rod-shaped UBE3B–CaM protomers are arranged in a closed, ring-like architecture, where the CaM subunit of each promoter acts as a molecular “glue” to mediate the head-to-tail dimeric assembly (Fig. 1, B to D). Within this dimeric structure, UBE3B features an N-terminal region (residues 1–58), comprising the N-terminal tail (N-tail; residues 1–28) and the IQ motif (residues 29–58), with the IQ motif serves as the CaM-binding site. Succeeding this helix is an Armadillo-like fold (residues 73–485), referred to as the Bridge domain. Protruding outward from the Bridge is a globular, triangular Hammer domain (residues 491–647). Finally, the C-terminus features the catalytic HECT domain (residues 650–1068), which consists of an N-lobe (residues 650–925) and a C-lobe (residues 933–1068). In our dimeric map, the C-lobe exhibits a very weak density due to its conformational heterogeneity, and is unresolved in the dimeric complex (Fig. 1C). However, in the trimer map, the Hammer domain of the third protomer stabilizes the HECT C-lobe of the adjacent subunit into a ‘T-shaped’ architecture while CaM acts as a fulcrum against the Bridge domain (fig. S2D). Taken together, these structural data represent snapshots of a propagating oligomerization process, providing structural evidence that UBE3B–CaM complexes are capable of forming higher-order assemblies.

**Fig. 2.**
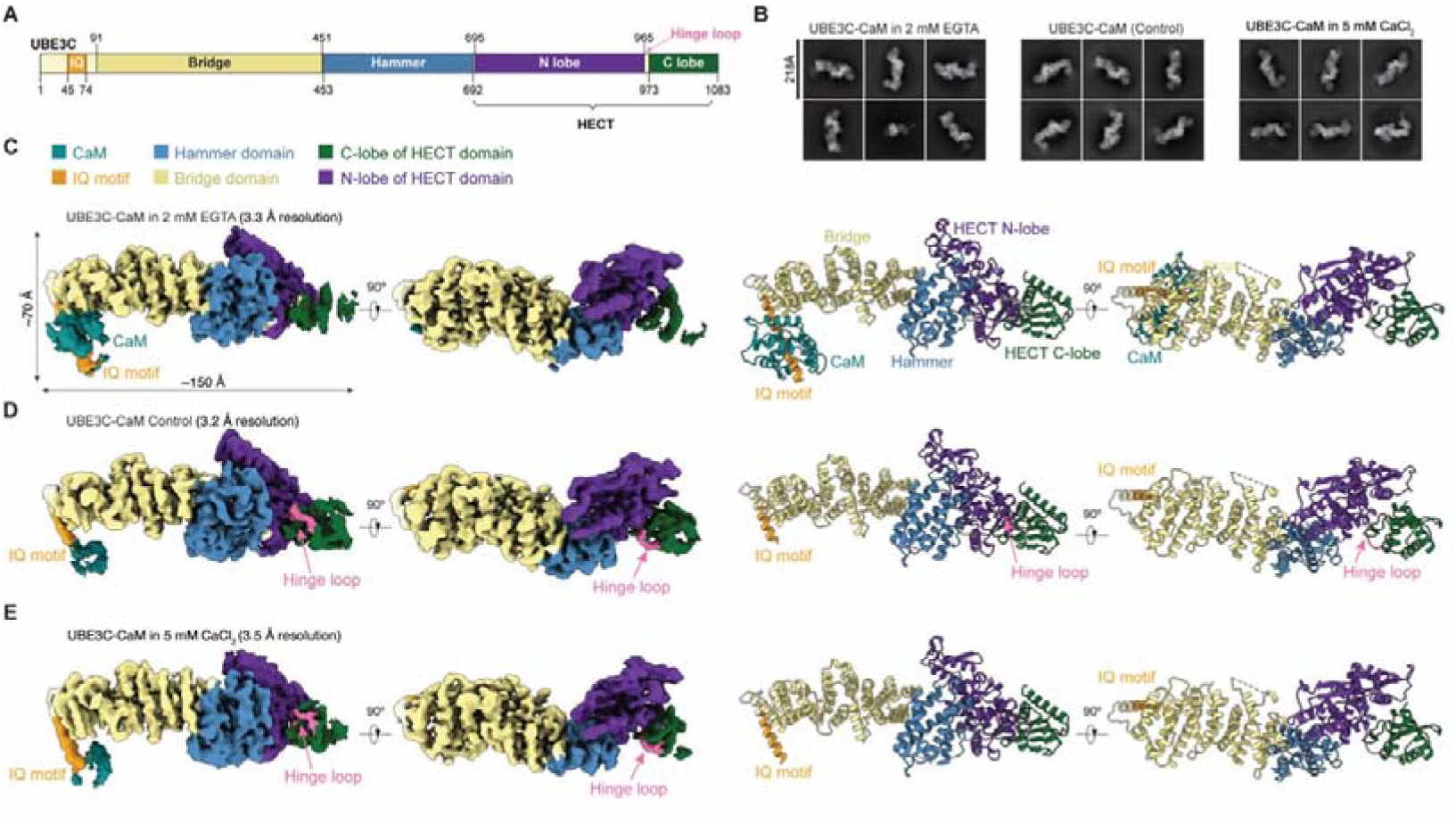
Structural landscapes of the UBE3C-CaM complex under various Ca^2+^ conditions. (**A**) Domain organization of human UBE3C, including the IQ motif, Bridge, Hammer, and HECT domain. (**B**) Representative 2D class averages of the UBE3C-CaM monomer in the presence of 2 mM EGTA (left), standard buffer (control) without added CaCl_2_ or EGTA (middle) and 5 mM CaCl_2_ (right). The box size is 218 Å for all conditions. (**C** to **E**) Cryo-EM density maps and corresponding structural models of UBE3C–CaM monomers obtained from datasets prepared under 2 mM EGTA (C) control (D) and 5mM CaCl_2_ (E) conditions, at overall resolutions of 3.3 Å, 3.2 Å, and 3.5 Å, respectively. Maps and models are color-coded by domain: IQ motif (orange), Bridge (yellow), Hammer (blue), HECT N-lobe (purple), HECT C-lobe (dark green), and CaM (dark cyan).

To investigate the structural impact of Ca^2+^, we also determined a cryo-EM map of the UBE3B–CaM complex in the presence of 5 mM CaCl_2_ at an overall resolution of 3.2 Å (Fig. 1F). Analysis of this dataset identified a predominantly monomeric population, with no significant higher-order classes detected (fig. S3A). While the core architecture of UBE3B remained largely consistent compared to the structure containing Ca^2+^-free CaM under the EGTA condition, Ca^2+^ binding to CaM significantly increased the inherent conformational flexibility of CaM and the UBE3B N-terminal region. This conformational plasticity was evidenced by diffuse density and reduced local resolution for these regions, which precluded de novo atomic model building for Ca^2+^-bound CaM (Fig. 1, B and E).

## Structures of human UBE3C–CaM complex

To uncover structural difference between UBE3C and UBE3B, we determined the cryo-EM structure of the UBE3C–CaM complex under three conditions: 2 mM EGTA (Fig. 2C), a standard buffer (control) without added CaCl_2_ or EGTA (Fig. 2D), and 5 mM CaCl_2_ (Fig. 2E), at overall resolutions of 3.3 Å, 3.2 Å and 3.5 Å, respectively. Representative 2D class averages confirmed CaM binding across all conditions (Fig. 2B). Analysis of these datasets did not identify any dimeric or oligomeric species, yielding only monomeric particles (fig. S3, B to D). The reconstructions reveal a compact, elongated L-shaped architecture, which maintains the conserved domain topology similar to UBE3B. The structure of UBE3C consists of an N-terminal IQ motif (residues 45–74), followed by the Bridge (residues 91–451), Hammer (residues 453–692), and the bilobed HECT (residues 695–1083) domains. The HECT domain comprises the N-lobe (residues 695–965) and C-lobe (residues 973–1083), connected by a flexible hinge loop (residues 966–972) (Fig. 2, B to E). The flexible N-tail (residues 1–44) is unresolvable in the map, indicating its highly flexible or disordered nature.

In the reconstruction of the EGTA dataset, CaM clearly binds to the N-terminal IQ motif of UBE3C (Fig. 2C). The global architecture of UBE3C remained largely consistent across the examined conditions, while the HECT domain exhibited variable dynamics. The control dataset yielded an improved local resolution for the catalytic HECT domain, where both the N-lobe and C-lobe are well resolved, and the catalytic cysteine (C1051) within the C-lobe is clearly defined (Fig. 2D; fig. S3D; fig. S5D).

Compared to the well-defined Ca^2+^-free CaM density observed in the EGTA condition, the CaM densities in both the control and 5 mM CaCl_2_ conditions appeared to be more diffuse and of lower local resolution, reflecting increased conformational dynamics of CaM. The flexibility observed in the control condition likely driven by trace amounts of endogenous Ca^2+^, suggesting that UBE3C-bound CaM is highly responsive to physiological calcium levels.

## CaM mediates UBE3B oligomerization

A striking feature of our cryo-EM structure is the central role of CaM in orchestrating the assembly of the UBE3B dimer. Structural analysis reveals that CaM bridges the two protomers, stabilizing the C2-symmetric assembly through distinct interfacial interactions (Fig. 1C). Specifically, the CaM C-lobe (residues 81–149) engages with the Bridge hairpin-loop (BHL) motif (residues 350–369) of the opposite protomer, burying a solvent-accessible surface area of 445.6 ± 77.4 Å^2^ (Fig. 3A). Structurally, the BHL motif comprises a short β1–β2 hairpin and a preceding ten-residue loop. This interaction is stabilized by a network of salt bridges formed between CaM residues D79, D81, E83, E84 and Bridge residues K352 and K353. Electrostatic surface analysis highlights charge complementarity across this interface, where the electronegative CaM C-lobe interacts with the electropositive BHL motif (Fig. 3B). In addition, the CaM N-lobe (residues 1-76) engages a positively charged region on the HECT domain of the opposite protomer, contributing a more modest interfacial area of 242.3 ± 27.2 Å^2^.

**Fig. 3.**
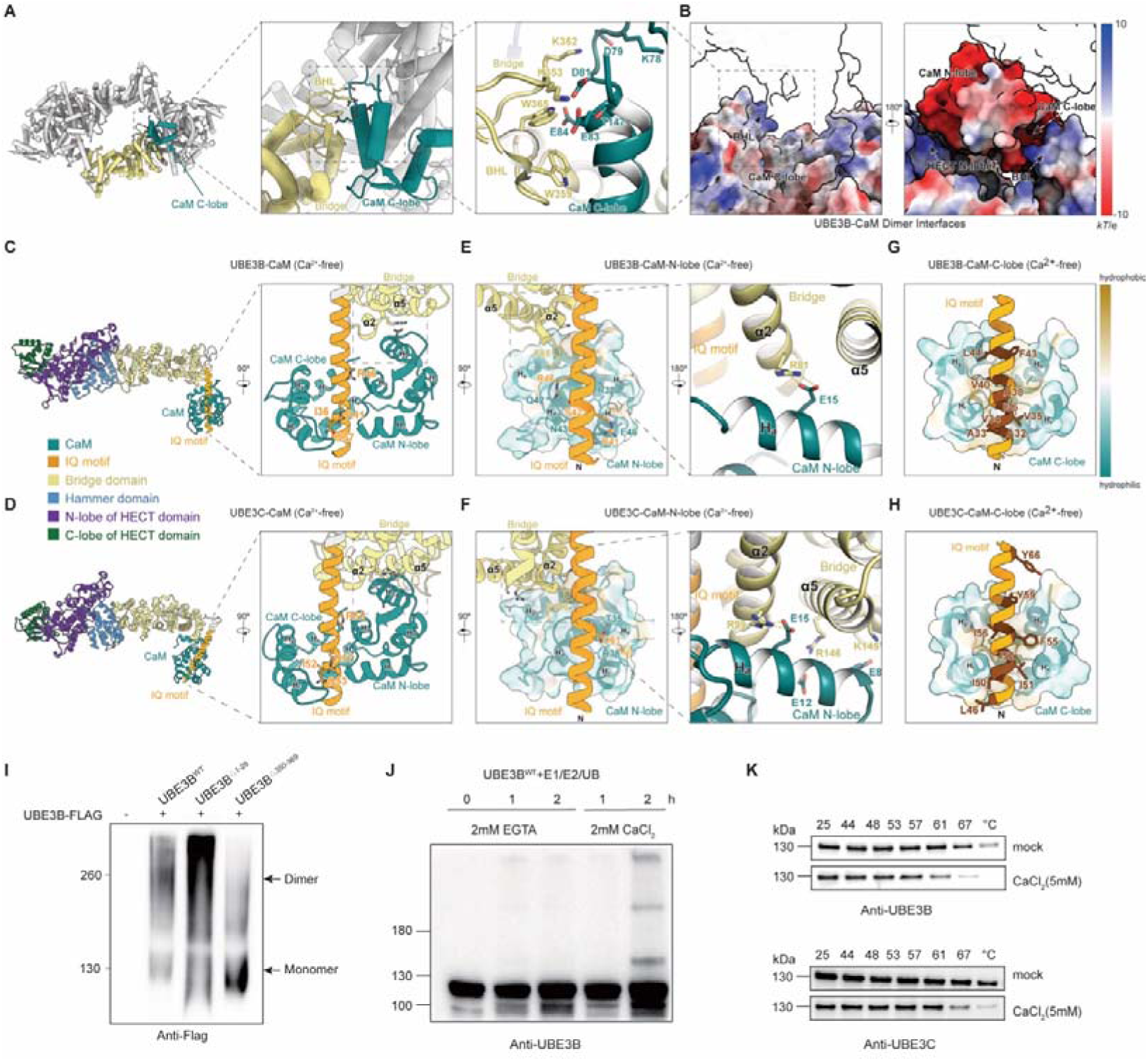
CaM-mediated stabilization of the UBE3B dimer and Ca^2+^-dependent regulation of UBE3B and UBE3C. **(A)** Overview (left) and detailed view (middle and right) of the UBE3B–CaM dimer interface showing contacts between CaM and the BHL motif. (**B**) Electrostatic surface of the UBE3B–CaM dimer interface involving BHL motif and HECT N-lobe, shown in two orientations rotated by 180° (blue, positive charge; red, negative charge). (**C**-**H)** Cryo-EM structures of UBE3B (C, E, and G) and UBE3C (**D**, **F**, and **H**) monomers in complex with Ca^2+^-free CaM (2 mM EGTA). The insets show the conserved residues of the IQ motif (C and D), the CaM N-lobe and the Bridge domain (E and F), and the CaM C-lobe-IQ motif hydrophobic interactions (G and H). (**I**) Native PAGE analysis of FLAG-tagged UBE3B^WT^ and mutants. (**J**) In vitro autoubiquitination of UBE3B^WT^ in the presence of 2 mM CaCl_2_ or 2 mM EGTA. (**K)** Cellular thermal shift assays of UBE3B and UBE3C in the presence or absence of CaCl_2_.

To investigate the role of the BHL motif in UBE3B dimerization, we analyzed wild-type (WT) and mutant UBE3B constructs expressed in HEK293T cells. Blue Native PAGE analysis of cell lysates confirmed that wild-type UBE3B forms a dimer under native conditions, whereas deleting the BHL motif (UBE3B^ΔBHL^)—which mediates the CaM interaction—markedly impaired dimerization (Fig. 3I). This strict dependence on CaM distinguishes UBE3B from other dimeric HECT ligases like UBR5 and HACE1, which oligomerize autonomously (fig. S1, H to J). By coupling dimerization to CaM recruitment, UBE3B acquires a unique regulatory mechanism that allows calcium signaling to modulate its assembly and ligase activity (fig. S1K).

## Ca^2+^-dependent regulation of UBE3B and UBE3C

Although CaM-mediated dimerization is unique to UBE3B, the fundamental recruitment of CaM via the IQ motif is conserved at both sequence and structure levels between UBE3B and UBE3C (Fig. 3, C to H; fig. S6, A to H). Our cryo-EM models reveal nearly identical binding modes for Ca^2+^-free CaM on the IQ motif of both ligases, burying total solvent-accessible surface area of 1623.6 Å^2^ in UBE3B and 1854.8 Å^2^ in UBE3C (Fig. 3, C and D; fig. S6, A and B). In both complexes, the IQ motif inserts deeply into the hydrophobic pocket of the CaM C-lobe, forming extensive interfaces of 936.7 Å^2^ in UBE3B and 1012.5 Å^2^ in UBE3C (Fig. 3, G and H). Structurally, the CaM C-lobe adopts a semi-open conformation while the N-lobe remains closed, a configuration consistent with the canonical Ca^2+^-free CaM binding mode observed in the MYO5A-IQ complex (*26*) (PDB 2IX7; fig. S6A).

Complementing the conserved C-lobe interaction, the Bridge domains of both UBE3B and UBE3C stabilize the CaM N-lobe through a shared anchoring mechanism (Fig. 3, E to F). UBE3B engages the CaM N-lobe primarily through a single α-helix (α2, residues 73–85), where Arg81 forms a salt bridge with CaM N-lobe residue Glu15, and Phe85 inserts into the CaM N-lobe hydrophobic pocket (Fig. 3E). UBE3C extends this interaction by employing two helices—an equivalent α-helix (α2, residues 91–104) and an additional helix (α5, residues 137–157)—to create a substantially larger interface (480.0 Å^2^) compared to UBE3B (304.7 Å^2^). Within this interface, Arg99, Lys145 and Arg146 form salt bridges with CaM N-lobe residues Glu15, Glu8 and Glu12, respectively. The conserved Phe103 similarly inserts into the hydrophobic pocket (Fig. 3F). These specific interactions effectively stabilize the CaM N-lobe, contrasting with the flexible conformation observed in the NMR structure of the Ca^2+^-free CaM–NaV1.5 IQ complex (*27*) (PDB 2L53; fig. S6C).

Structural superposition of the Ca^2+^-bound CaM–CAC1S IQ complex (*28*) (PDB 2VAY) onto the IQ motifs of our Ca^2+^-free CaM–UBE3B/C complexes reveals that Ca^2+^ binding induces an open conformation in the CaM N-lobe and C-lobe (fig. S6D). However, three-dimensional variability analysis (3DVA) reveals that, in the presence of Ca^2+^, CaM exhibits highly dynamic behavior in both UBE3B and UBE3C complexes, sampling a continuum of conformational states rather than a stable structure (fig. S6E). The increased conformational entropy is expected to disfavor the formation of higher-order assemblies, consistent with our cryo-EM observation of only monomeric form of UBE3B in the presence of Ca^2+^.

To assess the effect of Ca^2+^ on CaM binding, HEK293T cells were co-transfected with UBE3B^WT^ or UBE3C^WT^ together with CaM. The resulting complexes were immobilized on affinity beads and washed with buffers containing increasing concentrations of Ca^2+^. Immunoblot analysis revealed that CaM remains associated with both ligases across the Ca^2+^ titration range, indicating that Ca^2+^ binding does not appreciably dissociate CaM from UBE3B^WT^ and UBE3C^WT^ under these conditions (fig. S6, N and O). *In vitro* ubiquitylation assays demonstrated that Ca^2+^ facilitates the ubiquitin chain extension of both UBE3B^WT^ and UBE3C^WT^ (fig. S6, K and L).

Consistent with these findings, autoubiquitylation assays further indicated that Ca^2+^promotes the activity of both ligases (Fig. 3J; fig. S6P). To explore the underlying mechanism of Ca^2+^-dependent regulation, we purified the isolated HECT domains of UBE3B and UBE3C, designated as UBE3B^HECT^ (residues 700–1068) and UBE3C^HECT^(residues 693–1083). *In vitro* ubiquitylation assays revealed that Ca^2+^ had no obvious effect on the activity of these recombinant HECT domain proteins, suggesting that Ca^2+^ regulates the activity of UBE3B and UBE3C primarily through their N-terminal regions, consistent with our structural observations (fig. S6, I and J).

To further investigate how Ca^2+^-induced conformational dynamics impact the structural stability of the ligase complexes, we performed cellular thermal shift assays (CETSA). Lysates from HEK293T cells expressing the UBE3B or UBE3C complexes with CaM were subjected to a temperature gradient in the presence or absence of Ca^2+^, and the soluble fractions were analyzed by immunoblotting. The temperature-dependent assays revealed that the presence of 5 mM Ca^2+^ reduces the thermal stability of both UBE3B–CaM and UBE3C–CaM complexes compared to the Ca^2+^-free condition, as evidenced by their lowered melting temperatures (Fig. 3K).

Taken together, these observations suggest that Ca^2+^ binding to CaM orchestrates conformational dynamics of UBE3B/UBE3C-CaM complexes that facilitate their E3 ligase activity.

## Higher-order assembly-induced conformational remodeling

The HECT domains of UBE3B (residues 650–1068) and UBE3C (residues 695–1083) exhibit a high degree of sequence homology, sharing 44% sequence identity and 60% similarity (fig. S7). Consistent with the crystal structure of the truncated UBE3C HECT domain (PDB 6K2C) (*29*), our cryo-EM reconstruction of the full-length monomer displays a conserved L-shaped architecture, while also resolving flexible regions within the small subdomain of UBE3C HECT domain that were absent in the crystallographic model (Fig. 4F). While the L-shaped architecture represents the predominant conformation of the UBE3B monomer (Fig. 4E), 3DVA revealed pronounced flexibility within the catalytic HECT domains of both UBE3B and UBE3C monomers in the presence of 2 mM EGTA (fig. S8, A and B). Specifically, the rotation of the HECT domain relative to the Hammer domain generates a solvent-accessible cavity which accommodates the N-tail of the adjacent protomer during UBE3B dimerization (Fig. 4, A to C; fig. S8C). In the monomeric state of UBE3B, the N-terminal segment upstream of the IQ motif (residues 9–28) lacks stable density, indicating intrinsic disorder (Fig. 1E). This interaction buries a surface area of 859.2 ± 10.4 Å^2^ and is stabilized by hydrogen bonding and electrostatic complementarity between the positively charged N-tail and the negatively charged cavity (Fig. 4B).

**Fig. 4.**
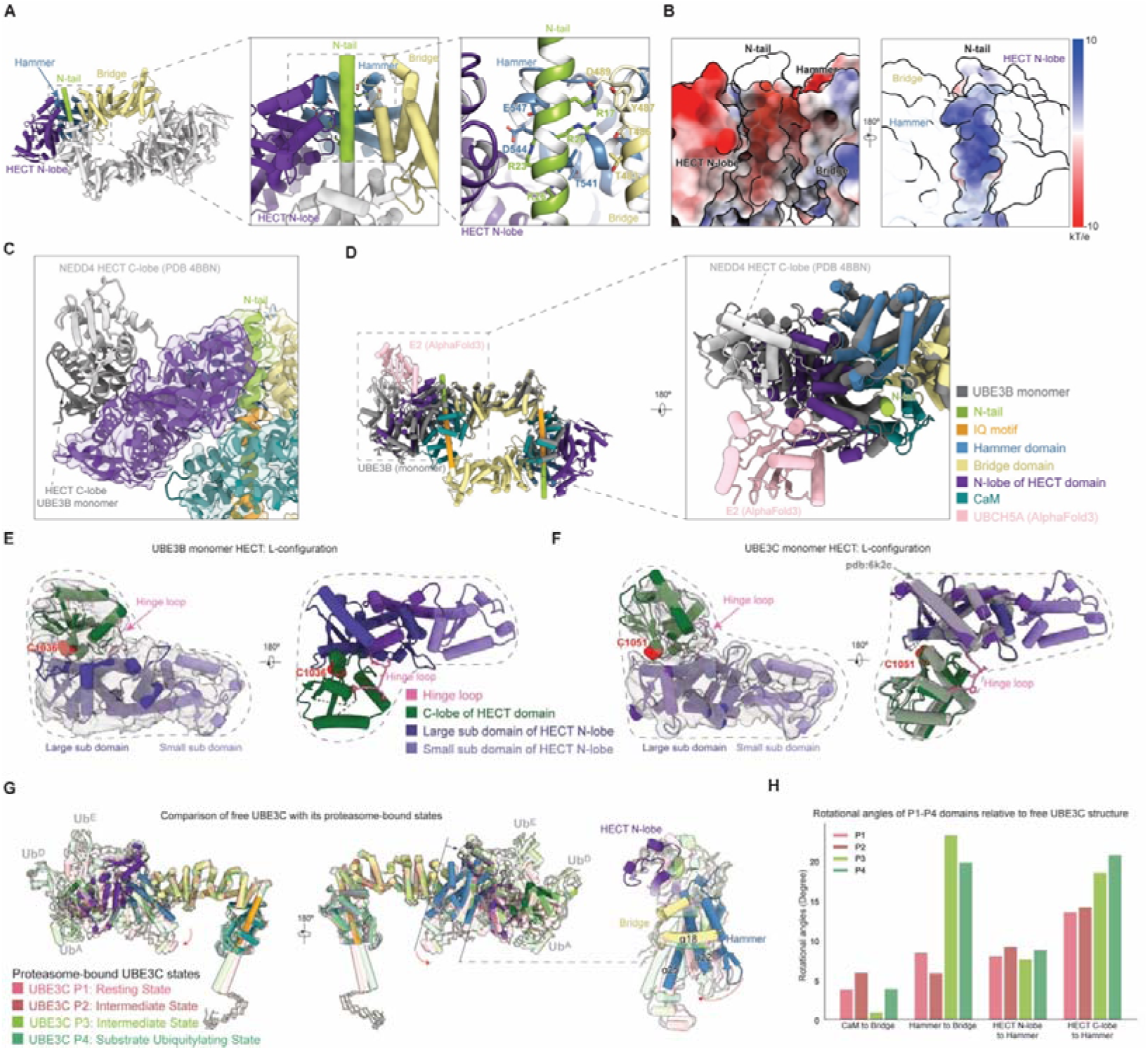
Cryo-EM structures and conformational dynamics of UBE3B and UBE3C. (**A**) Dimer interface showing the N-tail (green) insertion into the cavity of the adjacent protomer. Key interacting residues are labeled. (**B**) Electrostatic surface potential of the interacting surfaces, the negatively charged cavity (left, red) and the positively charged N-tail (right, blue). (**C**) Map (transparent surface) and atomic model of the dimer showing the N-tail (green) inserting into the dynamic cavity. Superimposed are the HECT C-lobes of the monomer (gray) and NEDD4 (light gray). (**D**) Superposition of an AF3-predicted UBE3B–CaM-UBCH5A (pink) on the experimental UBE3B–CaM dimer (aligned on the HECT N-lobe). The monomeric UBE3B (gray) is aligned on the Bridge domain for comparison. (**E** and **F**) Maps and atomic models of the L-shaped HECT domains of UBE3B (E) and UBE3C (F). Catalytic cysteines (C1036/C1051) and hinge loops, are indicated. The UBE3C HECT domain (PDB 6K2C, gray) is superimposed in (F). (**G**) Superposition of the proteasome-bound states (P1–P4) with the free UBE3C–CaM structure (solid, 2 mM EGTA state). (**H**) Quantification of inter-domain rotation angles of the proteasome-bound states (P1–P4) relative to the free UBE3C–CaM structure.

To investigate the conformational dynamics and stability differences between the monomeric and dimeric states of UBE3B, we performed 100-ns all-atom molecular dynamics (MD) simulations for both forms. The simulations confirmed that intermolecular interaction in the dimer significantly rigidify the N-tail, which is otherwise disordered in the monomer. This stabilization is evidenced by reductions in root mean square fluctuation (RMSF) values for the N-tail in the dimeric state (fig. S8D). Furthermore, the lower root mean square deviation (RMSD) observed in the dimer compared to the monomer supports the notion that dimerization locks flexible monomeric elements into a more rigid and stable assembly (fig. S8E). In agreement with the importance of the dimeric interface, cellular thermal shift assays showed that the dimerization-deficient mutant UBE3B^ΔBHL^ exhibits reduced stability, while deletion of the flexible N-tail (UBE3B^ΔN-tail^) had no significant effect (fig. S6M), underscoring that the BHL-mediated assembly core is the primary determinant of complex stability.

In a companion study, we report time-resolved cryo-EM reconstructions of the UBE3C–CaM–26S proteasome super-assembly in the act of ubiquitin chain elongation on a model substrate (*30*). Comparison with the free UBE3C structure determined here reveals that the proteasome-bound UBE3C adopts distinct conformations, characterized by markedly higher overall RMSD (3–5 Å) relative to the free state. This large conformational shift dictates allosteric regulation and activation by the 26S proteasome (fig. S8H). While individual domains (HECT N-lobe, C-lobe, Hammer, Bridge domains and CaM) remain rigid with low intra-domain RMSD values (fig. S8, F to H), the pronounced relative domain motions are observed, primarily between the Hammer and Bridge domains. Across the four captured conformational states (P1–P4) of proteasome-bound UBE3C, the Hammer domain undergoes a rotation of approximately 10–30° relative to the Bridge domain (Fig. 4, G and H). Consistently, a 500-ns all-atom MD simulation initiated from the P4-state (isolated from the proteasome), reproduces similar interdomain movements between the Hammer and Bridge. This simulation illustrates the relaxation trajectory of UBE3C as it transitions from its proteasome-bound conformation back to the free, equilibrated state (fig. S8, I and J). The inter-helical angles between the Hammer and Bridge domains converge toward values observed in simulations of the free UBE3C–CaM complex (fig. S8, J and L).

The N-terminal strand (residues 1–24) of UBE3C, which inserts into the lid subcomplex of the 26S proteasome, remains disordered in the absence of proteasome engagement. This behavior is analogous to that of the UBE3B N-tail, which requires dimerization for stabilization (Fig. 2, C to E; Fig. 4G). However, sequence and structural alignments show that the N-terminal strand responsible for 26S proteasome engagement is unique to UBE3C and is absent in UBE3B (fig. S7; fig. S10G).

## Modeling of substrate engagement and ubiquitin interactions

To assess the structural compatibility of the UBE3B dimer with ubiquitin ligase activity, we employed AF3 prediction and homology modeling to analyze the accessibility of key functional interfaces. Our analysis suggests that the dimeric conformation represents an active yet selectively constrained state, rather than a fully auto-inhibited one. Specifically, superposition of the AF3-predicted UBCH5A–UBE3B–CaM complex onto a single protomer of the UBE3B dimer reveals that the canonical E2-binding surface remains solvent-accessible, indicating that dimerization does not impose any steric hindrance on E2 recruitment (Fig. 4D; fig. S9C).

Furthermore, mapping the ubiquitin-binding sites observed in the proteasome-bound UBE3C structures (*30*) onto the UBE3B dimer reveals that the potential ubiquitin-binding surface on the HECT domain remains solvent-accessible (Fig. S9, A and B). This implies that the dimeric architecture appears structurally compatible with ubiquitin recruitment and the execution of ubiquitination reactions. However, substrate recognition appears to be hindered by dimerization according to AF3-predicted models of UBE3B–CaM monomer with various substrates, such as MYC (*11*), HIF-2α (*12*), PPP3CC (Protein Phosphatase 3 Catalytic Subunit Gamma) (*13*) and BCKDK (branched-chain α-keto acid dehydrogenase kinase) (*14*). By mapping the predicted per-residue contact probabilities onto the UBE3B–CaM dimer (Fig. 5a), we found that the primary substrate-recognition determinants cluster within the hand-like region formed by the Hammer domain with the adjacent HECT N-lobe and Bridge domain. This localization aligns well with previous truncation and mutagenesis studies on MYC and HIF-2α (Fig. 5B) (*11, 12*). Superposition of these AF3-predicted substrate complexes onto the dimer reveals that the substrate-binding sites spatially overlap with the dimeric interface (fig. S9, D to G). This suggests that dimerization acts as a conformational switch to restrict substrate access by occluding the recognition interface, thereby serving as a regulatory mechanism to tune its E3 activity toward specific targets. Similarly, mapping AF3-predicted contact probabilities for UBE3C substrates reveals that interaction hotspots are primarily concentrated on the HECT N-lobe, the Hammer–Bridge domains, and UBE3C-specific loop regions (Fig. 5C, fig. S9, H to J). This indicates that, beyond the catalytic HECT domain, the N-terminal regions are integral to the functional architecture of both UBE3B and UBE3C, presumably regulating substrate recruitment and specificity.

**Fig. 5.**
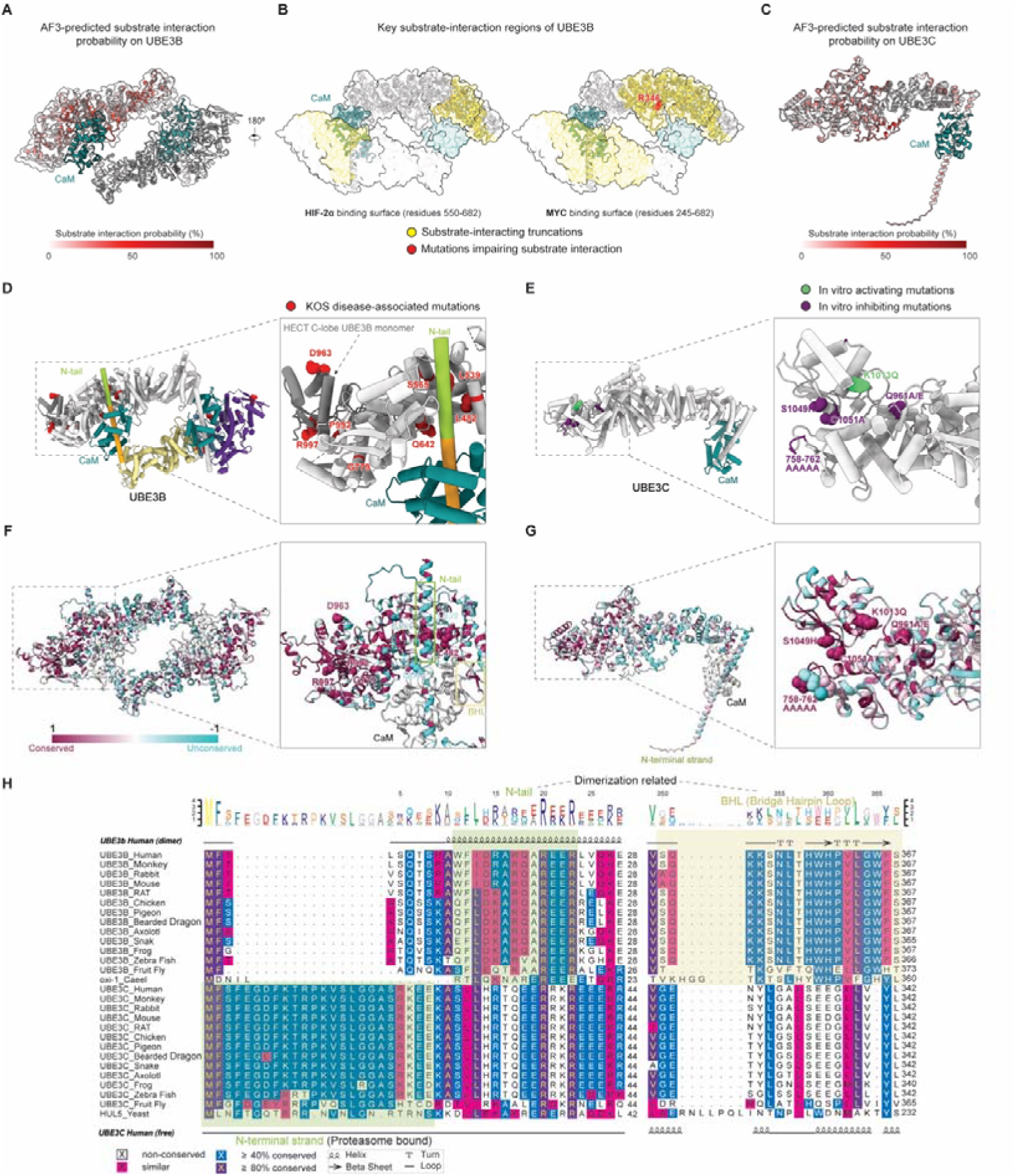
Substrate recognition, functional mapping and evolutionary divergence of UBE3B and UBE3C. (**A** and **C**) Surface representations of UBE3B (A) and UBE3C (C) colored by substrate interaction probability derived from AF3-predicted complexes (white, low; red, high). (**B**) Experimentally validated substrate-binding regions on UBE3B. Yellow surfaces: HIF-2α (residues 550–682) and MYC (residues 245–682) as previously reported; red spheres denote mutations impairing substrate ubiquitination. (**D**) Structural mapping of KOS-associated mutations (red spheres) onto the UBE3B–CaM dimer. The zoomed-in view highlights the clustering of pathogenic mutations involving the HECT domain and the N-tail-binding cavity on the Hammer domain. (**E**) Mapping of functionally characterized mutations onto the UBE3C structure. (green, activating; purple, inhibitory). (**F** and **G**) Evolutionary conservation scores mapped onto the structural models of the UBE3B dimer (**F**) and UBE3C (**G**) (cyan, variable; maroon, highly conserved). CaM is colored light gray. (**H**) Multiple sequence alignment (MSA) and secondary structures of UBE3B and UBE3C orthologs from representative species. Conserved residues are shaded based on similarity.

To determine the contribution of the non-catalytic N-terminal domains to catalytic efficiency, we compared the ubiquitin chain elongation activities of the isolated HECT domains (UBE3B^HECT^ and UBE3C^HECT^) with those of their full-length counterparts. Immunoblot analysis revealed that the E3 ligase activities of the HECT-only truncations were weaker than those of the full-length proteins (fig. S9, K and L). This suggests that the non-catalytic N-terminal domains are critical for potentiating the catalytic efficiency of ubiquitin chain elongation.

## Evolutionary divergence and functional specialization of UBE3B and UBE3C

Consistent with previous genomic studies (*31, 32*), a phylogenetic reconstruction places UBE3B and UBE3C in the Hul5-like clade of the UBE-subfamily, identifying yeast Hul5 as their common ancestor (fig. S10, A to C). Both human orthologs have acquired several loop insertions compared to Hul5 (Fig. S7A), and detailed multi-sequence alignment (MSA) across species further reveals their divergent architectural specialization (Fig. 5H; fig. S10, D and E). Specifically, UBE3C preserves the ancestral N-terminal strand (Fig. 5H, left), a motif critical for proteasome interaction, which is conserved across eukaryotes (Fig. 5G; fig. S10E). Within the catalytic HECT domain, the mutations known to experimentally modulate ligase activity (*29, 33*)—both inhibitory (purple) and activating (green)—are clustered around the catalytic cysteine C1051 (Fig. 5E). In contrast, UBE3B has lost this proteasome-binding strand but uniquely evolved the N-tail and the BHL motif (residues 350–369) (Fig. 5H, right). Mapping evolutionary conservation onto the UBE3B structure reveals that the dimerization interface—mediated by the N-tail, BHL motif, and the HECT–Hammer cavity—corresponds to a highly conserved surface patch (maroon), indicating strong evolutionary constraints on this assembly (Fig. 5F; fig. S10D). The high conservation of these UBE3B-specific elements from nematodes to mammals implies that the homodimeric architecture is an essential, evolutionarily selected feature of UBE3B function.

Mapping of pathogenic missense mutations associated with KOS syndrome cluster onto our UBE3B structure reveals their localization within both the catalytic HECT domain and the dynamic cavity (e.g., Leu482, Leu539) of the UBE3B dimer interface (*16, 34, 35*). While HECT domain mutations likely abolish catalytic turnover, interface mutations are predicted to compromise dimer formation (Fig. 5D). We speculate that such interface defects likely disrupt the Ca^2+^-sensing regulatory switch potentially by destabilizing and preventing proper resting-state assembly rather than simply causing a loss of intrinsic enzymatic activity.

Structural superposition of the UBE3C model onto the HECT domain of UBE3B dimer reveals that the distinct orientation of the UBE3C Bridge–CaM module would cause severe steric clashes with the Hammer domain of the adjacent protomer, precluding the formation of the same dimeric architecture observed in UBE3B (fig. S10F). Conversely, superimposing UBE3B onto the proteasome-bound UBE3C structure (aligned via the IQ–CaM module) reveals that, due to the distinct orientation of the Bridge–CaM module, the HECT–Hammer domain of UBE3B fails to engage with the proteasome lid, indicating that the UBE3B architecture is structurally incompatible with proteasome binding (fig. S10G). Furthermore, the evolution of several additional flexible loops alongside the preserved N-terminal strand, likely provides the necessary conformational plasticity to support optimal catalytic function upon proteasome recruitment (fig. S7; fig. S8L). The differential retention and evolution of these structural elements constitute the molecular basis for their distinct higher-order assemblies: a CaM-dependent dimerization interface for UBE3B, and the conserved N-terminal strand for proteasome recruitment in UBE3C.

## Discussion

Higher-order assembly has emerged as essential regulatory mechanisms for diverse enzymatic systems. Several HECT E3 ligase family members, such as E6AP (*36–38*), UBR5 (*39–42*) and HACE1 (*43, 44*), assemble as dimers or trimers that are critical for regulating their enzymatic activities. Our cryo-EM structures of CaM-mediated oligomeric architectures of UBE3B reveal a previously unrecognized regulatory paradigm in which E3 superassemblies are controlled by calcium signaling (Fig. 6). To our knowledge, this study provides the first structural elucidation of how Ca^2+^ signaling modulates higher-order superassemblies of E3 ubiquitin ligases via CaM acting as an inter-protomer “molecular glue”. Notably, this mode of CaM-mediated regulation and the asymmetric trimer formed on an anti-parallel homodimer in the UBE3B superassemblies are distinct from all previously reported E3 oligomeric architectures (*36–47*). In contrast to prior E3 dimerization mechanisms that rely on intrinsic structural elements (e.g., the N-terminal helix of HACE1 (*43, 44*), HECT–HECT stacking in HUWE1 (*47*), or ARM repeats in UBR5 (*39–42*)), UBE3B assembly is governed by an extrinsic Ca^2+^ sensor, CaM, which is sandwiched between adjacent E3 protomers and confers pronounced Ca^2+^ sensitivity in both ligase activities and structural dynamics (fig. S1, H to K).

**Fig. 6.**
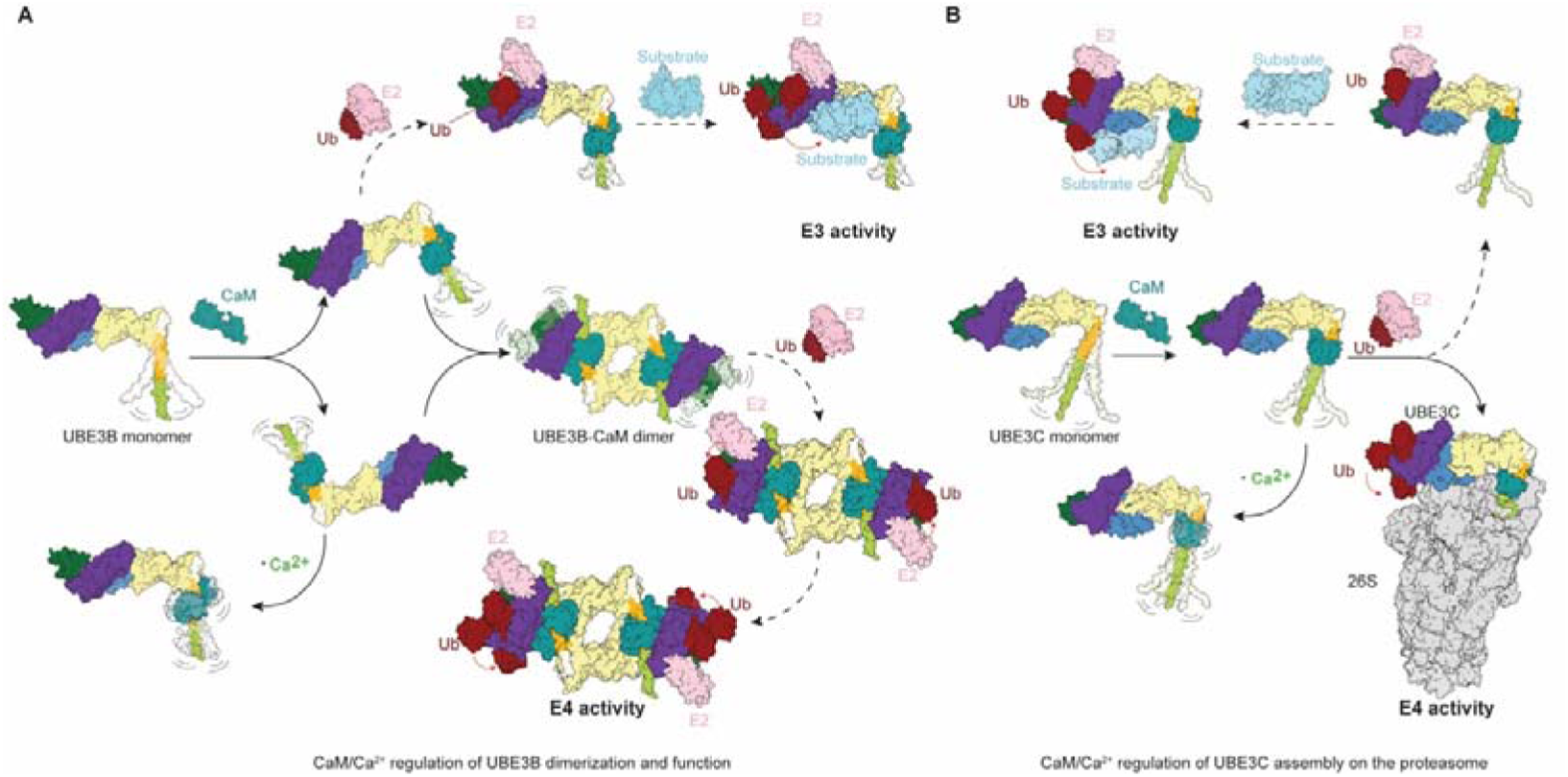
Mechanistic hypothesis of CaM/Ca^2+^-dependent regulation of UBE3B and UBE3C. (**A**) A proposed mechanistic model illustrating the CaM-mediated modulation of UBE3B dimerization and its E3 vs potential E4 activities. In low Ca^2+^, CaM binding facilitates the assembly of the UBE3B homodimer. The monomeric form primarily functions as an E3 ligase for substrate recognition and initiation, whereas the homodimer is hypothesized to regulate the E3 ligase specificity and ubiquitin chain elongation activity (E4 activity). Ca^2+^ influx induces conformational dynamics in CaM, promoting dimer dissociation and potentially recycling the ligase back to the monomeric state. (**B**) A proposed mechanistic model illustrating CaM-mediated modulation of UBE3C and its E3 vs E4 activities. The UBE3C–CaM monomer functions as an E3 ligase for substrate recognition and initiation. Upon association with the 26S proteasome via its flexible N-terminal strand, UBE3C is primed for enhanced processive E4 activity on proteasome-bound substrates. Ca^2+^ binding introduces structural plasticity that may regulate the dwell time or facilitate the release of UBE3C from the proteasome. General Note: Solid lines represent experimentally determined structures resolved by cryo-EM, while dashed arrows indicate structures derived from AF3 prediction or homology modeling.

By integrating AF3 predictions and homology modelling with our structural analyses, we demonstrate that the higher-order assembly exhibits distinct spatial selectivity. Of note, the oligomeric configuration is compatible with ubiquitin chain elongation, maintaining an E4 activity (fig. S9B). However, superposition with substrate-bound models indicates that oligomerization partially occludes the substrate-recognition interfaces in several cases (Fig. 5A; fig. S9, D to G). Thus, oligomerization may act as a key regulatory mechanism that toggles UBE3B between E3 and E4 activities, by spatially modulating substrate accessibility (Fig. 5A; fig. S10H). This mechanism underscores the functional plasticity of HECT family ligases and helps explain how they meet the multifaceted demands of protein quality control.

UBE3C exhibits E3 activity in isolation and shows pronounced E4 activity when associating with the proteasome (*30*). To enable proteasome recruitment, UBE3C utilizes its N-terminal strand (absent in UBE3B) and has evolved several unique loops (Fig. 5H; fig. S7A). These structural adaptation is reflected in the enhanced mobility observed between the Hammer and Bridge domains, as well as the distinct Bridge–CaM orientation of UBE3C compared to UBE3B (Fig. 4, G and H; fig. S10, C and D). CaM-modulated orientation of UBE3C N-terminal region is therefore critical for efficient proteasome association (*30*). Furthermore, Ca^2+^ influx may promote CaM conformational flexibility to facilitate UBE3C dissociation from the proteasome, thereby switching its E4 activity back to its E3 ligase activity (Fig. 6B) (*30*).

Physiologically, UBE3B localizes to the outer mitochondrial membrane (*9*), where resting cytosolic Ca^2+^ is tightly maintained at a basal level of approximately 100 nM (*48*). We speculate that this low Ca^2+^ concentration environment favors the formation of substrate-selective higher-order assemblies. Compared to the monomer, the UBE3B dimer exhibits increased flexibility of the HECT C-lobe. Notably, in the trimeric assembly, one protomer functions as an allosteric driver that biases the C-lobe of the adjacent protomer toward a T-shaped conformation poised for efficient ubiquitin transfer from the E2 enzyme. Taken together, our findings support a model in which the higher-order assembly adopts a primed state, ready for rapid ubiquitin transfer and optimized for responsiveness to upstream signaling cues (fig. S10H). Elevation in intracellular Ca^2+^—triggered by synaptic transmission (*49*) or ROS-induced mitochondrial release (*50*)—serve as a molecular switch through Ca^2+^ binding to UBE3B/C-associated CaM, altering structural dynamics of UBE3B/C-CaM and promoting disassembly of the higher-order complexes. This Ca^2+^-driven transition from oligomers into active monomers is predicted to generate an ultrasensitive, cooperative and switch-like response to Ca^2+^ signals, converting UBE3B/C from E4 to E3 activities (Fig. 6; fig. S10H). By preserving an E2∼Ub–compatible geometry in the catalytic domain after higher-order assembly, this mechanism may enable rapid deployment of the enzymes for the high-throughput targeted ubiquitylation of stress-responsive substrates, providing an essential layer of Ca^2+^ signaling-dependent regulation of the ubiquitin-proteasome system and autophagy.

## Materials and Methods

### Constructs

Full-length human *UBE3B* cDNA was synthesized (Kangwei Biotech) and subcloned into a modified pcDNA3.1 vector containing an N-terminal 6×His–biotin tag followed by a Tobacco Etch Virus (TEV) protease cleavage site. Mutant constructs of *UBE3B*, including the N-terminal deletion (Δ1–28), the isolated HECT domain (residues 700–1068) and the BHL deletion (Δ350–369), were generated on the same vector backbone. For cellular thermal shift (CETSA) and native state analyses (Blue Native PAGE, affinity precipitation), *UBE3B* and *UBE3C* variants were subcloned into the pCDH-CMV-MCS-EF1-Puro vector containing Flag tags. All constructs were verified by Sanger sequencing (Qingke Biotech).

Similarly, full-length human *UBE3C* was cloned into a modified pcDNA3.1 vector engineered to include an N-terminal 6×His–biotin tag and a TEV protease cleavage site. The UBE3C HECT domain (residues 693–1083) was subcloned into the pGEX-4T-1 vector as an N-terminal GST-fusion protein. Full-length human calmodulin 1 (*CaM*) was subcloned into the pcDNA3.1 vector for transient expression without affinity tags. The cDNA encoding human UBE1 was a gift from C. Tang, and the human UBCH5A cDNA was purchased from GenScript; both were subcloned into pGEX-4T-1 for expression as N-terminal GST-fusion proteins. Recombinant human ubiquitin was obtained from Boston Biochem.

### Cell culture

The HEK293T cell line was obtained from the National BioMedical Cell Resource Center (BMCR, China), and the Expi293 cell line was a gift from J. Sodroski. All cell lines tested negative for mycoplasma contamination. HEK293T cells were cultured in Dulbecco’s Modified Eagle’s Medium (DMEM; Gibco) supplemented with 10% fetal bovine serum (FBS; Gibco) and 1% penicillin–streptomycin at 37 °C in a humidified incubator containing 5% CO_2_. Expi293 cells were maintained in SMM 293-TII medium (SinoBiological) under the same conditions in suspension culture.

### Protein expression and purification

Full-length human UBE3B and UBE3C were expressed in Expi293F cells. For each preparation, 200 ml of Expi293F culture at a density of 2 × 10^6^ cells/ml was co-transfected with *UBE3B* and *CaM* (or *UBE3C* and *CaM*) plasmids at a 1:1 mass ratio. A total of 360 µg plasmid DNA was mixed with 1080 µg polyethylenimine (PEI; Yeasen) in Opti-MEM (Gibco) for 15 min before transfection. After 12 h, 10 mM sodium butyrate was added to enhance expression, and cells were harvested 48 h post-transfection. Cell pellets were lysed in buffer containing 50 mM Tris–HCl pH 7.5, 150 mM NaCl, 10% glycerol, 2 mM ethylene glycol-bis(β-aminoethyl ether)-N,N,N′,N′-tetraacetic acid (EGTA), 1 mM tris(2-carboxyethyl)phosphine (TCEP), 0.5% NP-40, supplemented with protease inhibitors (Roche). Cells were lysed by sonication on ice, and the lysate was clarified by centrifugation at 20,000□g for 30□min at 4□°C.

For UBE3B purification, the supernatant was incubated with Ni–Sepharose resin (Yeasen) for 3h at 4 °C. The resin was washed with 20 column volumes (CV) of lysis buffer, followed by 10 CV of wash buffer (50 mM Tris–HCl pH 7.5, 150 mM NaCl, 10% glycerol, 1 mM TCEP, and 10 mM imidazole). Bound UBE3B–CaM complexes were eluted with elution buffer (50 mM Tris–HCl pH 7.5, 150 mM NaCl, 10% glycerol, 1 mM TCEP, and 300 mM imidazole). Eluted proteins were digested with 15 µl of TEV protease (2 mg/ml; laboratory stock, plasmid was a gift from Y. Yin) overnight at 4 °C. TEV-cleaved proteins were further purified by size-exclusion chromatography (SEC) using a Superdex 200 Increase 10/300 GL column (Cytiva) equilibrated in SEC buffer (25 mM Tris–HCl pH 7.5, 100 mM NaCl, 1 mM TCEP, and 2 mM EGTA).

For UBE3C purification, the supernatant was incubated with Streptavidin Beads 6FF (Smart-Lifesciences) for 3 h at 4 °C. The resin was washed with 20 CV of lysis buffer, followed by 10 CV of wash buffer (50 mM Tris–HCl pH 7.5, 150 mM NaCl, 10% glycerol, and 1 mM TCEP). Bound UBE3C–CaM complexes were released by on-column TEV cleavage in cleavage buffer (50 mM Tris–HCl pH 7.5, 100 mM NaCl, and 1 mM TCEP) overnight at 4 °C. The eluted proteins were purified by SEC using SEC buffer (25 mM Tris–HCl pH 7.5, 100 mM NaCl, 1 mM TCEP, 2 mM EGTA).

Human UBE1 and UBCH5A were expressed as N-terminal GST-fusion proteins in *E. coli* BL21-CodonPlus (DE3)-RIPL cells. Bacterial cultures were grown to an OD_600_ of 0.6–0.7 and induced with 0.2 mM isopropyl β-D-1-thiogalactopyranoside (IPTG) (20 h at 16 °C for UBE1; overnight at 18°C for UBCH5A). Cells were harvested and lysed by sonication in lysis buffer containing 25 mM Tris–HCl (pH 7.5), 150 mM NaCl, 10 mM MgCl_2_, 0.2% Triton X-100, 1 mM dithiothreitol (DTT), and protease inhibitors. The supernatant was incubated with Glutathione Sepharose 4B resin (Yeasen) for 3 h at 4 °C, washed extensively with wash buffer (25 mM Tris–HCl pH 7.5, 150 mM NaCl, and 1 mM DTT), and digested on-column with thrombin (Solarbio) in cleavage buffer (20 mM Tris–HCl pH 8.0, 150 mM NaCl, and 1 mM DTT) overnight at 4 °C to remove the GST tag. UBE1 was further purified using a Superdex 200 Increase 10/300 GL column (Cytiva), and UBCH5A was purified using a Superdex 75 Increase 10/300 GL column (Cytiva), both equilibrated in buffer containing 25 mM Tris–HCl (pH 7.5), 150 mM NaCl, 1 mM DTT, and 10% glycerol.

### Ca2+-dependent interaction assay

Expi293F cells were co-transfected with pcDNA3.1(+) vectors encoding His–biotin–UBE3B or His–biotin–UBE3C and CaM. Cells were harvested and lysed in a buffer containing 50 mM Tris–HCl (pH 7.5), 150 mM NaCl, 10% glycerol, 2 mM EGTA, 1 mM TCEP, 0.5% NP-40, and 1× protease inhibitor cocktail. Lysates were clarified by centrifugation (20,000 × *g*, 30 min, 4 °C) and incubated with Streptavidin Agarose Resin 6FF for 2h at 4 °C. The beads were washed with lysis buffer and evenly divided into aliquots. Each aliquot was resuspended in a base buffer (50 mM Tris–HCl pH 7.5, 150 mM NaCl, 1 mM TCEP) supplemented with 0, 1, 5, or 25 mM CaCl[. The mixtures were incubated at 4 °C for 10 min with gentle agitation to allow equilibrium binding. Subsequently, the beads were washed three times with the same base buffer (50 mM Tris–HCl pH 7.5, 150 mM NaCl, 1 mM TCEP) supplemented with 0, 1, 5, or 25 mM CaCl_2_, respectively, to maintain the specific calcium concentration of each group. Bound proteins were eluted with 1× SDS loading buffer and analyzed by SDS–PAGE and immunoblotting using anti-CaM (Abcam, 1:2000 dilution) and anti-UBE3B (Abmart, 1:2000) or anti-UBE3C (Abcam, 1:2000) antibodies.

### *In vitro* ubiquitylation assays

To reconstitute the E3 ligase activity of UBE3B and UBE3C, *in vitro* ubiquitylation assays were performed in a reaction buffer containing 50 mM HEPES (pH 7.5), 100 mM NaCl, 10 mM MgCl_2_, 5 mM ATP, and 1 mM TCEP. All reactions contained 0.15 µM recombinant human UBE1, 5 µM UBCH5A, and 30 µM ubiquitin.

For solution-based assays (ubiquitin chain elongation), purified full-length or HECT-domain truncated UBE3B/UBE3C proteins (1 µM) were incubated with the reaction mixture (containing 50 mM HEPES pH 7.5, 100 mM NaCl, 10 mM MgCl_2_, 1 mM TCEP, 5 mM ATP, 0.15 µM UBE1, 5 µM UBCH5A, and 30 µM ubiquitin) at 37 °C for the indicated time periods. To examine Ca^2+^-dependent regulation, CaCl_2_ or EGTA was added to the specified final concentrations. Reactions were terminated by the addition of 5× SDS loading buffer and boiling at 95 °C for 5 min. Reaction products were resolved by SDS–PAGE and analyzed by immunoblotting using linkage-specific anti-polyubiquitin antibodies (ABclonal, 1:2000 dilution).

For on-bead assays (auto-ubiquitylation), Expi293F cells transiently transfected with pcDNA3.1 vectors encoding His–biotin–UBE3B or UBE3C were lysed in a buffer containing 50 mM HEPES (pH 7.5), 150 mM NaCl, 10% glycerol, 2 mM EGTA, 0.5% NP-40, and 1× protease inhibitor cocktail. Lysates were clarified by centrifugation (20,000 × *g*, 20 min, 4 °C) and incubated with Streptavidin Agarose Resin 6FF for 2h at 4 °C. The beads were washed extensively with lysis buffer and equilibrated in the reaction buffer. Bead-immobilized complexes were then resuspended in 10–20 µL of the aforementioned reaction mixture supplemented with the indicated concentrations of CaCl_2_ or EGTA and incubated at 37 °C. Reactions were stopped by boiling in SDS loading buffer, and ubiquitylation signals were detected by Western blotting using anti-UBE3B (Abmart, 1:2000 dilution) or anti-UBE3C (Abcam, 1:2000 dilution) antibodies.

### Blue Native PAGE

HEK293T cells transfected with pCDH vectors encoding UBE3B or UBE3C were harvested, washed with ice-cold phosphate-buffered saline (PBS), and lysed in BN-lysis buffer containing 25 mM HEPES (pH 7.0), 25 mM NaCl, 10% glycerol, 2 mM EGTA, 0.1% NP-40, and 1× protease inhibitor cocktail for 30 min on ice. Lysates were clarified by centrifugation at 20,000 × *g* for 20 min at 4 °C. The supernatants (20 µg of total protein) were mixed 1:1 (v/v) with 2× BN-sample buffer (100 mM Bis-Tris HCl, 500 mM 6-aminocaproic acid, 30% glycerol, 5% Serva Blue G) and resolved on 4–13% gradient Blue Native PAGE gels (BeyoGel). Electrophoresis was performed at 4 °C, initially at 110 V for 30 min using Cathode Buffer A (50 mM Tricine, 7.5 mM imidazole, 0.02% Serva Blue G, pH 7.0), followed by 250 V for 1 h using Cathode Buffer B (50 mM Tricine, 7.5 mM imidazole, pH 7.0, 0.002% Serva Blue G). Proteins were transferred to polyvinylidene fluoride (PVDF) membranes (300 mA for 90 min at 4 °C) and analyzed by immunoblotting as described in the Immunoblotting section.

### Cellular thermo-stability assay

HEK293T cells (2×10^5^ per well, 6-well plates) were co-transfected with CaM and either pCDH –UBE3C, UBE3B(WT), or UBE3B variants (Flag-tagged mutants). Cells were washed with PBS, lysed in 500 µL buffer (50 mM HEPES pH 7.5, 150 mM NaCl, 10% glycerol, 2 mM EGTA, 0.5% NP-40, and 1× protease inhibitor cocktail) for 30 min on ice, and centrifuged (20,000 × g, 20 min, 4 °C). Aliquots (50 µL) were heated for 15 min at the indicated temperatures (25–67 °C), cooled on ice for 5 min, and centrifuged again (15,000 × *g* for 20 min at 4 °C). Supernatants (20 µL) were mixed with 5 µL 5× SDS loading buffer and analyzed by SDS–PAGE and immunoblotting with anti-UBE3B or anti-UBE3C antibodies.

### BS3 crosslinking

Purified UBE3B–CaM complexes were buffer-exchanged by desalting into 25 mM HEPES pH 7.5 and 150 mM NaCl, and adjusted to 1 mg/ml.

Bis(sulfosuccinimidyl)suberate (BS3) was freshly dissolved in ice-cold water to 50 mM immediately before use and added to the protein sample to a final concentration of 1–2 mM. Reactions were incubated at 37 °C for 30 min. After incubation, crosslinking reactions were quenched by adding Tris–HCl (pH 7.5) to a final concentration of 50 mM and incubated at room temperature for 15 min, followed by SDS loading buffer and denaturation at 95 °C for 5 min. Samples were analyzed by SDS–PAGE and Western blot using anti-UBE3B and anti-CaM antibodies.

### Mass photometry

Mass photometry assays were conducted using the TwoMP mass photometer (Refeyn, UK). Microscope coverslips and CultureWell gaskets were purchased from Refeyn.

The gaskets were placed on cleaned coverslips on the sample stage of the mass photometer, following the manufacturer’s instructions. BS3-crosslinked UBE3B–CaM and UBE3C–CaM complexes were freshly purified and buffer-exchanged into MP buffer (20 mM HEPES pH 7.4, 150 mM NaCl) using Zeba spin desalting columns (Thermo Fisher Scientific). The MP buffer was freshly filtered through a 0.22 µm filter prior to measurement. After dilution in MP buffer to a final concentration of approximately 20 nM, 10 µL of protein samples were placed in the sealed wells, and light scattering was recorded for 60 s. Data calibration was performed using a standard protein mixture containing markers of known molecular weights (66 kDa, 224 kDa, and 660 kDa) to generate a standard curve. Data were acquired using AcquireMP (Refeyn) and analyzed with DiscoverMP (Refeyn), followed by Gaussian fitting to determine the molecular masses of the monomeric (∼125 kDa), dimeric (∼250 kDa), and trimeric (∼389 kDa) species.

### Cryo-EM sample preparation and data collection

Purified UBE3B–CaM and UBE3C–CaM complexes were buffer-exchanged into imaging buffer (50 mM Tris–HCl pH 6.8, 100 mM NaCl, 1 mM DTT) using 7 kDa desalting columns and adjusted to 0.8 mg/ml. After buffer exchange, samples were supplemented with the indicated Ca^2+^ or EGTA concentrations and incubated at 25 °C for 30 min. For UBE3B–CaM, cryo-EM grids were prepared under three biochemical conditions: 2 mM EGTA (used for both a monomer dataset and a dimer dataset, with the latter additionally collected at 20°, 30° and 40° stage tilt), and 5 mM CaCl_2_. For UBE3C–CaM, samples were prepared in parallel under three conditions: 2 mM EGTA, 5 mM CaCl_2_, or without the addition of either CaCl_2_ or EGTA following buffer exchange. Cryo-EM grids were prepared using an FEI Vitrobot Mark IV (Thermo Fisher Scientific). All specimens were vitrified on glow-discharged Quantifoil R1.2/1.3 Au 300 mesh grids (H□/O□ plasma, 50 W, 80 s) at 4 °C and 100% humidity with blot force −2 and blot time 2 s, followed by plunge-freezing into liquid ethane.

Cryo-grids were initially screened using a 200 kV Tecnai Arctica microscope (Thermo Fisher Scientific). High-quality grids were transferred to a 300 kV Titan Krios G2 microscope (Thermo Fisher Scientific), equipped with a post-column BioQuantum energy filter (Gatan) connected to a K3 direct electron detector (Gatan). Coma-free alignment and parallel illumination were manually optimized before each data collection session. Cryo-EM data were automatically acquired using SerialEM software (*51*) in super-resolution counting mode with 20 eV energy slit. All datasets were collected at a calibrated super-resolution pixel size of 0.425 Å, with a nominal defocus range of -1.4 to -1.8 µm and a total accumulated dose of ∼60 electrons per Å^2^.

For the UBE3B datasets, 8,751 movies were collected in 2 mM EGTA (untilted), and 5,789 movies were collected for the Ca^2+^-bound state (5 mM Ca^2+^). To ameliorate the preferred orientation issue observed in the UBE3B dimer sample (2 mM EGTA), additional data were collected with stage tilts, yielding 2011 movies at 20°, 6,179 movies at 30° and 4,871 movies at 40°. For UBE3C, datasets were collected as follows: 3,084 movies for Ca^2+^-free state (2 mM EGTA), 3,205 movies for Ca^2+^-bound state (5 mM Ca^2+^), and 5,975 movies for control (without additional EGTA or Ca^2+^).

### Cryo-EM data processing

For UBE3B and UBE3C monomer in the presence of 2 mM EGTA or 5mM Ca^2+^, beam-induced motion correction was performed using the MotionCor2 (*52*) implementation within RELION (*53*) version 4.0 and 5.0, with a super-resolution pixel size of 0.425 Å. Contrast transfer function (CTF) parameters of the drift-corrected micrographs were calculated using the Gctf program (*54*). Micrographs were subsequently curated in cryoSPARC (*55*) v4.7.1 to remove those exhibiting crystalline ice, cracks, significant astigmatism, or poor CTF fit resolution (>5 Å). Initial particle picking was performed in cryoSPARC via template matching, using 2D projections derived from AF3 predicted models. These models were first converted into 3D density maps in UCSF ChimeraX (*56*) and low-pass filtered to 20 Å before projection. Subsequent 2D and 3D classification with fourfold binned particles yielded a clean dataset suitable for further analysis. Topaz (*57*) was trained on the cleaned subset and used to re-pick particles in cryoSPARC to maximize recovery. Then auto-refinement with application of the Blush algorithm (*58*), CTF refinement and Bayesian polishing, was carried out in RELION using two-fold binned particles at a pixel size of 0.85 Å. For UBE3C in 2 mM EGTA, subsequent 3D classification with skip-alignment in RELION and AlphaCryo4D (*59*) analysis yielded a clean dataset of 545,496 particles, resulting in a final reconstruction at 3.3 Å resolution that revealed improved density for the HECT domain. For UBE3B monomer (2 mM EGTA), UBE3B (5 mM Ca^2+^), UBE3C (5 mM Ca^2+^) and UBE3C (without additional EGTA or Ca^2+^), the final datasets contain 689,636, 369,410, 486,419 and 789,678 particles, resulting in a final reconstruction at 2.9, 3.2, 3.5 and 3.2 Å, respectively. 3D variability analysis (3DVA) (*60*) was performed in cryoSPARC using the final particle stacks with a filter resolution of 8 Å and solving for the top 3 principal variability modes (number of modes = 3).

For the UBE3B dimer dataset in the presence of 2 mM EGTA, beam-induced motion correction and CTF estimation were performed using Patch Motion and Patch CTF implementations in cryoSPARC. Initial particle picking was carried out on the untilted micrographs using Blob Picking. These particles were used to generate an initial 3D reference via Ab-Initio Reconstruction. 2D projections from this selected volume were then used for template-based picking across the entire dataset, including the untilted and 20°, 30° and 40° tilted micrographs. Following 2D classification and Heterogeneous Refinement with fourfold binned particles to remove damaged or incomplete particles, Topaz training and picking were used to maximize the recovery of high-resolution and rare orientation particles, yielding an original dataset of 2,232,151 particles. To overcome the severe preferred orientation problem during reconstruction, we employed a combined strategy using CryoPROS (*61*), spIsoNet (*62*) and CryoSieve (*63*). The original dataset (2,232,151 particles) and a low-pass filtered map from a previous refinement served as the initial latent volume for CryoPROS training, after which auxiliary particles were generated to match the size of the original dataset but with a balanced angular distribution. This expanded particle stack (doubled in size) was subjected to co-refinement using cryoSPARC Non-Uniform refinement. Following 2D classification to down-weight overrepresented views, the resulting sharpened map from the intrinsic dataset was used as the latent volume for a second round of CryoPROS training and generation. This particle expansion strategy amplified the clustering features of minor orientations, thereby improving the selection accuracy for high-tilt views. A balanced subset was then curated by recycling orientationally rebalanced particles, resulting in a final intrinsic set of 900,876 particles. These particles were re-extracted using two-fold binned particles at a pixel size of 0.85 Å and subjected to auto-refinement with C1 symmetry in RELION, integrated with spIsoNet to correct for misalignment caused by preferred orientation. During this process, 10 epochs of inference were performed in each local search iteration. The refined set was subsequently optimized using CryoSieve based on the established alignments and half-set splits. To minimize overfitting, a low-pass filter of 4.5 Å was applied during the RELION postprocessing step within the iterative sieving. After 7 rounds of sieving with a retention ratio of 0.8, a final subset of 188,926 particles was retained. This high-quality subset yielded a final reconstruction at 3.3 Å resolution.

During the classification of the UBE3B–CaM dataset (2 mM EGTA), we observed distinct 2D class averages indicative of a trimeric assembly (Extended Data Fig. 2c). To investigate this structure, we pooled all curated micrographs from the untilted and tilted (0°–40°) datasets, totaling 19,878 micrographs. Particles were initially selected based on the specific trimeric 2D classes and used to generate an initial 3D model via Ab-Initio Reconstruction. Projections of this volume were then used for template picking, followed by Topaz training and picking to maximize the enrichment of this subpopulation. A final set of 221,081 trimeric particles was subjected to Non-Uniform Refinement in cryoSPARC and local refinement integrated with spIsoNet in RELION, yielding a reconstruction at 4.0 Å resolution. To visualize the molecular architecture, the atomic models of the UBE3B–CaM dimer and monomer were independently docked into the map using rigid-body fitting. The resulting assembly reveals a conformation where the HECT C-lobe mediating the interaction with the third (“upper”) UBE3B subunit displays relatively well-defined density, adopting a T-shaped conformation (fig. S2D).

### Structural analysis and visualization

Initial structural models for all monomers were generated using AlphaFold3 (AF3) predictions of the full complexes. These models were docked into the cryo-EM density maps, manually adjusted in Coot (*64*) and subsequently optimized by real-space refinement in PHENIX (*65*). For the UBE3B dimer, two copies of the AF3-predicted UBE3B–CaM model were independently fitted into the dimer cryo-EM map as rigid bodies, followed by the same protocol of atomic model refinement. In the final atomic models, highly flexible regions with poor density were truncated. For UBE3B structures, a large internal loop in the Bridge domain (residues 405–451) is consistently disordered and absent in all models. The N-terminal disordered region varies slightly among the states: residues 1–9 were removed in the dimer, whereas residues 1–28 and 1–38 were truncated in the apo monomer and Ca^2+^-bound monomer, respectively. In the UBE3B dimer map, the HECT C-lobe (residues 926–1068) exhibited weak density due to significant conformational flexibility. Given the ambiguity of its precise orientation, the entire C-lobe and two additional loops (residues 672–678 and 717–726) were considered unresolved and excluded from the final atomic coordinates. The monomeric UBE3B structures (both Ca^2+^-free and Ca^2+^-bound) retain the C-lobe, with only a specific loop (residues 999–1031) within the lobe being unresolved. For UBE3C structures, four internal loops (residues 360–384, 524–532, 562–591, and 633–676) are consistently unresolved across all conditions. The N-terminal truncation varies, with residues 1–44, 1–61, and 1–57 missing in the EGTA, control, and Ca^2+^-bound datasets, respectively. Regarding CaM, the first three N-terminal residues are missing in all structures solved under EGTA condition, while the entire CaM molecule is unresolved in the UBE3B (5 mM CaCl_2_) or UBE3C (control, 5 mM CaCl_2_) structure.

Interface analysis was performed using the PDBePISA server (*66*) (EMBL-EBI). For the UBE3B dimer, the interface area was calculated by isolating the specific interacting pairs from the C2-symmetric assembly. The solvent-accessible surface areas are reported as the average of the values calculated for the two symmetric interfaces, with the difference between them indicated as the uncertainty. Cryo-EM maps and atomic models were analyzed using Coot, PyMOL (*67*), UCSF Chimera (*68*), and ChimeraX (*56*). Global resolution was estimated based on the gold-standard Fourier Shell Correlation (FSC) = 0.143 criterion using the SPIDER (*69*) software package. Map-to-model FSC curves were calculated using PHENIX (FSC = 0.5 criterion). Angular distribution plots and directional FSC analysis for the UBE3B dimer were analyzed in cryoSPARC. Local resolution was estimated using ResMap (*65*) and visualized in UCFS Chimera. All structural figures preparation and alignments for visualization were prepared using UCSF ChimeraX and PyMOL.

### Molecular dynamics simulation

Molecular dynamics (MD) simulations were performed using the GROMACS 2022 package (*70*) with the Amber14SB force field. Simulation systems for the Ca^2+^-free UBE3B–CaM dimer, UBE3B–CaM monomer, and UBE3C–CaM monomer were constructed based on the cryo-EM structures solved in this study, with missing loops and flexible regions supplemented by AF3 predictions. The Hul5–CaM complex was modeled directly with AF3. Each system was solvated with TIP3P water molecules in a triclinic box, with a minimum distance of 1.5 nm maintained between the protein surface and the box edges. The systems were neutralized by the addition of appropriate counterions (Na^+^ or Cl^−^). Energy minimization was performed using the conjugate gradient algorithm until the maximum force converged to less than 100 kJ mol^-1^ nm^-1^. The systems were then equilibrated in the NPT ensemble (298.15 K, 1 bar) for 100 ps. During equilibration, temperature was controlled with the v-rescale thermostat and pressure with the Berendsen barostat, and positional restraints were applied to all protein heavy atoms. Production MD simulations were performed for 100 ns per system in the NPT ensemble (298.15 K, 1 bar) with a 2-fs time step. For the production runs, the pressure coupling was switched to the Parrinello-Rahman barostat. Long-range electrostatic interactions were handled with the Particle Mesh Ewald (PME) method, and all bonds involving hydrogens were constrained. Trajectories were recorded every 2 ps. For analysis, the first 40 ns were discarded as equilibration. Root-mean-square deviation (RMSD) and root-mean-square fluctuation (RMSF) were calculated over the subsequent stable segment. To simulate the transition of UBE3C from the P4 state to the free state, an extended production run of 500 ns was performed following the same protocol. The time evolution of the inter-helical angles for UEB3C Hammer–Bridge conformational changing is analyzed in fig. S8J, L. To calculate these angles, the axis of each helix (Bridge: α18, residues 420–435; Hammer: α22, 490–512 and α25, 565–571) was defined as the vector connecting the center of mass of its six N-terminal residues to that of its six C-terminal residues.

### Homology modeling and AlphaFold 3 predictions

Homology modeling of the UBE3B (HECT)–ubiquitin complex was performed using the Modeller (*71*) program with UCSF ChimeraX (*56*), utilizing the P1-P4 proteasome-bound UBE3C (HECT)–ubiquitin structures (*30*) as templates. All structural predictions were performed using the AlphaFold 3 web server (https://alphafoldserver.com/) (*25*). To identify substrate-interaction hotspots (Fig. 5a–b), we implemented a contact probability analysis based on multiple independent predictions. For UBE3B–substrate interactions, we performed 20 independent predictions with four distinct random seeds for each substrate (e.g., MYC (*11*), HIF2α (*12*), PPP3CC (*13*), BCKDK (*14*)). For UBE3C–substrate interactions, a similar protocol was applied using 10 independent predictions (two random seeds generating five models each) for a panel of substrates including ANXA7 (Annexin-7) (*72*), cyclin B1 (*73*), DOT1L (DOT1 Like Histone Lysine Methyltransferase) (*74*), PEBP1 (Phosphatidylethanolamine Binding Protein 1) (*19*), TP73 (Tumor Protein P73) (*75*), VPS34 (Vacuolar protein sorting 34) (*76*), and AXIN1 (*18*). An interaction site was defined as any UBE3B/C residue with at least one atom located within 5 Å of the substrate in a given model. For each residue, contact probability was calculated as the frequency of being classified as an interaction site across all predicted models. These probability values were mapped onto the UBE3B–CaM dimer or UBE3C–CaM structures and visualized using UCSF ChimeraX. The confidence of the predicted models was evaluated based on the predicted Local Distance Difference Test (pLDDT) scores and Predicted Aligned Error (PAE) matrices. The per-residue pLDDT scores were mapped onto the 3D structures using UCSF ChimeraX, and the PAE matrices were visualized using Python scripts.

### Multiple sequence alignment and structure alignment

Multiple sequence alignments (MSA) was generated using the MAFFT algorithm (*77*) or the CLUSTALW algorithm (*78*) with the “SLOW/ACCURATE” setting to maximum precision. The resulting alignments were visualized and annotated using the ESPript 3.0 server (*79*) and TEXshade software (*80*). For structural comparisons, the TM-align algorithm (*81*) was employed to calculate the Root-Mean-Square Deviation (RMSD) and rotation matrices between the free UBE3C structure against its various domain conformations bound to the proteasome (P1-P4 states). The rotation angles presented in Fig. 4H and fig. S8H were derived directly from these calculated rotation matrices.

### Phylogenetic analysis of HECT E3 ligases

To examine the evolutionary relationship between Hul5 and vertebrate UBE3B/UBE3C, we constructed a phylogenetic tree based on HECT domain sequences across representative species. The Pfam (*82*) HECT-domain HMM (PF00632) was used to search proteomes from fungi, protists, invertebrates, and vertebrates using HMMER hmmsearch (*83*) with an E-value cutoff of 1e^−5^ and a minimum domain coverage of 60%. For each hit, the HECT region was extracted according to the ali-from/ali-to coordinates reported by hmmsearch, and redundant or highly similar isoforms were removed to generate a non-redundant sequence set.

Multiple sequence alignment was performed using MAFFT L-INS-I (*77*), followed by trimming of poorly aligned regions with TrimAl (automated1 mode) (*84*).

Maximum-likelihood phylogenetic trees were generated using IQ-TREE 2 (*85*) with automatic model selection and 1,000 ultrafast bootstrap (*86*) and 1,000 SH-aLRT replicates (*87*). The final Newick tree was visualized and annotated in iTOL (*88*), with clades manually highlighted to distinguish the yeast Hul5 lineage, invertebrate UBE3 family members, and vertebrate UBE3B/UBE3C clades, thereby revealing the evolutionary divergence between Hul5 and its metazoan orthologs.

## Acknowledgements

The cryo-EM data were collected at the Cryo-EM Core Facility Platform and Laboratory of Electron Microscopy at Peking University. The data processing was supported by Biomedical Computing Platform of National Biomedical Imaging Center, the High-Performance Computing Platform, and the Weiming No. 1 and Life Science No. 1 supercomputing systems, at Peking University. This work was supported in part by National Natural Science Foundation of China (12125401, 12090051 and 32471308), National Key Research and Development Program of China (2023YFF1204400 and 2023YFF1204401), Beijing Natural Science Foundation grant (Z180016/Z18J008) and by AI for Science (AI4S)-Preferred Program at Peking University Shenzhen Graduate School.

## Contributions

X.L., J.H. and S.Z. carried out protein preparation, *in vitro* assays, cell-based assays and cryo-EM sample preparation. M.S., X.L., J.H., M.L., G.W. and Y.H. performed cryo-EM data collection. M.S., Q.W. and D.Y. processed and analyzed the cryo-EM data. M.S. conducted MD simulation, homology modelling, AF3 prediction, MSA analysis, and structural visualizations. X.L. performed phylogenetic analysis. Y.M., X.L. and M.S. wrote the manuscript. Y.M. designed the study and supervised the project, with input from all authors.

## Data availability

The cryo-EM density maps and corresponding atomic coordinates determined in this study have been deposited in the Electron Microscopy Data Bank (EMDB) and Protein Data Bank (PDB), respectively. All other data supporting the findings of this study are available from the corresponding author upon reasonable request.

## Notes

### Competing Interest Statement

The authors have declared no competing interest.

## REFERENCES

1. I. Dikic, Proteasomal and Autophagic Degradation Systems. Annu Rev Biochem 86, 193–224 (2017).

2. C. Pohl, I. Dikic, Cellular quality control by the ubiquitin-proteasome system and autophagy. Science 366, 818–822 (2019).

3. M. Lazarou et al., The ubiquitin kinase PINK1 recruits autophagy receptors to induce mitophagy. Nature 524, 309–314 (2015).

4. N. Zheng et al., Structure of the Cul1-Rbx1-Skp1-F boxSkp2 SCF ubiquitin ligase complex. Nature 416, 703–709 (2002).

5. M. G. Koliopoulos, D. Esposito, E. Christodoulou, I. A. Taylor, K. Rittinger, Functional role of TRIM E3 ligase oligomerization and regulation of catalytic activity. EMBO J 35, 1204–1218 (2016).

6. V. Balaji, T. Hoppe, Regulation of E3 ubiquitin ligases by homotypic and heterotypic assembly. F1000Res 9, 88 (2020).

7. J. J. Work, O. Brandman, Adaptability of the ubiquitin-proteasome system to proteolytic and folding stressors. J Cell Biol 220, e201912041 (2021).

8. B. W. Chu et al., The E3 ubiquitin ligase UBE3C enhances proteasome processivity by ubiquitinating partially proteolyzed substrates. J Biol Chem 288, 34575–34587 (2013).

9. A. Braganza et al., UBE3B Is a Calmodulin-regulated, Mitochondrion-associated E3 Ubiquitin Ligase. J Biol Chem 292, 2470–2484 (2017).

10. Y. Gu et al., Nrf2/UBE3B protects against acute lung injury by inhibiting ferritinophagy through the ubiquitination of NCOA4. Biol Direct 20, 85 (2025).

11. K. Li et al., TRIB3 promotes MYC-associated lymphoma development through suppression of UBE3B-mediated MYC degradation. Nat Commun 11, 6316 (2020).

12. Y. Wang et al., UBE3B promotes breast cancer progression by antagonizing HIF-2alpha degradation. Oncogene 42, 3394–3406 (2023).

13. M. C. Ambrozkiewicz et al., The murine ortholog of Kaufman oculocerebrofacial syndrome protein Ube3b regulates synapse number by ubiquitinating Ppp3cc. Mol Psychiatry 26, 1980–1995 (2021).

14. S. Cheon et al., The ubiquitin ligase UBE3B, disrupted in intellectual disability and absent speech, regulates metabolic pathways by targeting BCKDK. Proc Natl Acad Sci U S A 116, 3662–3667 (2019).

15. C. Sun et al., UBE3C tunes autophagy via ATG4B ubiquitination. Autophagy 20, 645–658 (2024).

16. E. Flex et al., Loss of function of the E3 ubiquitin-protein ligase UBE3B causes Kaufman oculocerebrofacial syndrome. J Med Genet 50, 493–499 (2013).

17. A. N. Cutrupi et al., Novel gene-intergenic fusion involving ubiquitin E3 ligase UBE3C causes distal hereditary motor neuropathy. Brain 146, 880–897 (2023).

18. Y. Zhang et al., UBE3C promotes proliferation and inhibits apoptosis by activating the beta-catenin signaling via degradation of AXIN1 in gastric cancer. Carcinogenesis 42, 285–293 (2021).

19. Z. Xu et al., Circular RNA circPOLR2A promotes clear cell renal cell carcinoma progression by facilitating the UBE3C-induced ubiquitination of PEBP1 and, thereby, activating the ERK signaling pathway. Mol Cancer 21, 146 (2022).

20. C. Hang, S. Zhao, T. Wang, Y. Zhang, Oncogenic UBE3C promotes breast cancer progression by activating Wnt/beta-catenin signaling. Cancer Cell Int 21, 25 (2021).

21. E. M. Kawamoto, C. Vivar, S. Camandola, Physiology and pathology of calcium signaling in the brain. Front Pharmacol 3, 61 (2012).

22. M. Zhang et al., Structural basis for calmodulin as a dynamic calcium sensor. Structure 20, 911–923 (2012).

23. M. Bhattacharyya et al., Molecular mechanism of activation-triggered subunit exchange in Ca(2+)/calmodulin-dependent protein kinase II. Elife 5, (2016).

24. X. Li et al., Calmodulin dissociates the STIM1-Orai1 complex and STIM1 oligomers. Nat Commun 8, 1042 (2017).

25. J. Abramson et al., Accurate structure prediction of biomolecular interactions with AlphaFold 3. Nature 630, 493–500 (2024).

26. A. Houdusse et al., Crystal structure of apo-calmodulin bound to the first two IQ motifs of myosin V reveals essential recognition features. Proc Natl Acad Sci U S A 103, 19326–19331 (2006).

27. B. Chagot, W. J. Chazin, Solution NMR structure of Apo-calmodulin in complex with the IQ motif of human cardiac sodium channel NaV1.5. J Mol Biol 406, 106–119 (2011).

28. D. B. Halling et al., Determinants in CaV1 channels that regulate the Ca2+ sensitivity of bound calmodulin. J Biol Chem 284, 20041–20051 (2009).

29. S. Singh, J. Sivaraman, Crystal structure of HECT domain of UBE3C E3 ligase and its ubiquitination activity. Biochem J 477, 905–923 (2020).

30. S. Zou et al., UBE3C retrofits the proteasome to enforce degradation of ultra-stable folds. bioRxiv, (2026).

31. I. Marin, Origin and evolution of fungal HECT ubiquitin ligases. Sci Rep 8, 6419 (2018).

32. M. C. Ambrozkiewicz, K. J. Cuthill, D. Harnett, H. Kawabe, V. Tarabykin, Molecular Evolution, Neurodevelopmental Roles and Clinical Significance of HECT-Type UBE3 E3 Ubiquitin Ligases. Cells 9, 2455 (2020).

33. E. A. Faqeih et al., Biallelic variants in HECT E3 paralogs, HECTD4 and UBE3C, encoding ubiquitin ligases cause neurodevelopmental disorders that overlap with Angelman syndrome. Genet Med 25, 100323 (2023).

34. L. Basel-Vanagaite et al., Expanding the clinical and mutational spectrum of Kaufman oculocerebrofacial syndrome with biallelic UBE3B mutations. Hum Genet 133, 939–949 (2014).

35. A. Albakheet et al., Novel UBE3B mutations: report of eight patients with Kaufman oculocerebrofacial syndrome with additional clinical findings from a highly consanguineous population. Clin Dysmorphol 33, 55–62 (2024).

36. L. Huang et al., Structure of an E6AP-UbcH7 complex: insights into ubiquitination by the E2-E3 enzyme cascade. Science 286, 1321–1326 (1999).

37. Z. Wang et al., Structural insights into the functional mechanism of the ubiquitin ligase E6AP. Nat Commun 15, 3531 (2024).

38. V. P. Ronchi, J. M. Klein, D. J. Edwards, A. L. Haas, The active form of E6-associated protein (E6AP)/UBE3A ubiquitin ligase is an oligomer. J Biol Chem 289, 1033–1048 (2014).

39. L. A. Hehl et al., Structural snapshots along K48-linked ubiquitin chain formation by the HECT E3 UBR5. Nat Chem Biol 20, 190–200 (2024).

40. F. Wang et al., Structure of the human UBR5 E3 ubiquitin ligase. Structure 31, 541–552 e544 (2023).

41. J. M. Tsai et al., UBR5 forms ligand-dependent complexes on chromatin to regulate nuclear hormone receptor stability. Mol Cell 83, 2753–2767 e2710 (2023).

42. Z. Hodakova et al., Cryo-EM structure of the chain-elongating E3 ubiquitin ligase UBR5. EMBO J 42, e113348 (2023).

43. S. Singh et al., Structural Basis for the Enzymatic Activity of the HACE1 HECT-Type E3 Ligase Through N-Terminal Helix Dimerization. Adv Sci (Weinh) 10, e2207672 (2023).

44. J. During et al., Structural mechanisms of autoinhibition and substrate recognition by the ubiquitin ligase HACE1. Nat Struct Mol Biol 31, 364–377 (2024).

45. Z. Yang et al., Molecular basis of SIFI activity in the integrated stress response. Nature 643, 1117–1126 (2025).

46. D. B. Grabarczyk et al., Architecture of the UBR4 complex, a giant E4 ligase central to eukaryotic protein quality control. Science 389, 909–914 (2025).

47. M. Hunkeler et al., Solenoid architecture of HUWE1 contributes to ligase activity and substrate recognition. Mol Cell 81, 3468–3480 e3467 (2021).

48. M. J. Berridge, P. Lipp, M. D. Bootman, The versatility and universality of calcium signalling. Nat Rev Mol Cell Biol 1, 11–21 (2000).

49. M. J. Berridge, The Inositol Trisphosphate/Calcium Signaling Pathway in Health and Disease. Physiol Rev 96, 1261–1296 (2016).

50. A. Gorlach, K. Bertram, S. Hudecova, O. Krizanova, Calcium and ROS: A mutual interplay. Redox Biol 6, 260–271 (2015).

51. D. N. Mastronarde, Automated electron microscope tomography using robust prediction of specimen movements. J Struct Biol 152, 36–51 (2005).

52. S. Q. Zheng et al., MotionCor2: anisotropic correction of beam-induced motion for improved cryo-electron microscopy. Nat Methods 14, 331–332 (2017).

53. J. Zivanov et al., New tools for automated high-resolution cryo-EM structure determination in RELION-3. Elife 7, (2018).

54. K. Zhang, Gctf: Real-time CTF determination and correction. J Struct Biol 193, 1–12 (2016).

55. A. Punjani, J. L. Rubinstein, D. J. Fleet, M. A. Brubaker, cryoSPARC: algorithms for rapid unsupervised cryo-EM structure determination. Nat Methods 14, 290–296 (2017).

56. T. D. Goddard et al., UCSF ChimeraX: Meeting modern challenges in visualization and analysis. Protein Sci 27, 14–25 (2018).

57. T. Bepler et al., Positive-unlabeled convolutional neural networks for particle picking in cryo-electron micrographs. Nat Methods 16, 1153–1160 (2019).

58. D. Kimanius et al., Data-driven regularization lowers the size barrier of cryo-EM structure determination. Nat Methods 21, 1216–1221 (2024).

59. Z. Wu, E. Chen, S. Zhang, Y. Ma, Y. Mao, Visualizing Conformational Space of Functional Biomolecular Complexes by Deep Manifold Learning. Int J Mol Sci 23, 8872 (2022).

60. A. Punjani, D. J. Fleet, 3D variability analysis: Resolving continuous flexibility and discrete heterogeneity from single particle cryo-EM. J Struct Biol 213, 107702 (2021).

61. H. Zhang et al., CryoPROS: Correcting misalignment caused by preferred orientation using AI-generated auxiliary particles. Nat Commun 16, 4565 (2025).

62. Y. T. Liu, H. Fan, J. J. Hu, Z. H. Zhou, Overcoming the preferred-orientation problem in cryo-EM with self-supervised deep learning. Nat Methods 22, 113–123 (2025).

63. J. Zhu et al., A minority of final stacks yields superior amplitude in single-particle cryo-EM. Nat Commun 14, 7822 (2023).

64. P. Emsley, K. Cowtan, Coot: model-building tools for molecular graphics. Acta Crystallogr D Biol Crystallogr 60, 2126–2132 (2004).

65. A. Kucukelbir, F. J. Sigworth, H. D. Tagare, Quantifying the local resolution of cryo-EM density maps. Nat Methods 11, 63–65 (2014).

66. E. Krissinel, K. Henrick, Inference of macromolecular assemblies from crystalline state. J Mol Biol 372, 774–797 (2007).

67. Schrödinger LLC, The PyMOL Molecular Graphics System, Version 2.4.0. (Schrödinger, LLC, New York, 2020).

68. E. F. Pettersen et al., UCSF Chimera--a visualization system for exploratory research and analysis. J Comput Chem 25, 1605–1612 (2004).

69. T. R. Shaikh et al., SPIDER image processing for single-particle reconstruction of biological macromolecules from electron micrographs. Nat Protoc 3, 1941–1974 (2008).

70. M. J. Abraham et al., GROMACS: High performance molecular simulations through multi-level parallelism from laptops to supercomputers. SoftwareX 1**-****2**, 19–25 (2015).

71. B. Webb, A. Sali, Comparative Protein Structure Modeling Using MODELLER. Curr Protoc Bioinformatics 54, 5 6 1-5 6 37 (2016).

72. S. J. Pan et al., Ubiquitin-protein ligase E3C promotes glioma progression by mediating the ubiquitination and degrading of Annexin A7. Sci Rep 5, 11066 (2015).

73. M. Okada et al., Liganded ERalpha Stimulates the E3 Ubiquitin Ligase Activity of UBE3C to Facilitate Cell Proliferation. Mol Endocrinol 29, 1646–1657 (2015).

74. T. Song et al., DOT1L O-GlcNAcylation promotes its protein stability and MLL-fusion leukemia cell proliferation. Cell Rep 36, 109739 (2021).

75. Y. Wang, M. Liu, X. Liu, X. Guo, LINC00963-FOSB-mediated transcription activation of UBE3C enhances radioresistance of breast cancer cells by inducing ubiquitination-dependent protein degradation of TP73. J Transl Med 21, 321 (2023).

76. Y. H. Chen et al., VPS34 K29/K48 branched ubiquitination governed by UBE3C and TRABID regulates autophagy, proteostasis and liver metabolism. Nat Commun 12, 1322 (2021).

77. K. Katoh, D. M. Standley, MAFFT multiple sequence alignment software version 7: improvements in performance and usability. Mol Biol Evol 30, 772–780 (2013).

78. J. D. Thompson, D. G. Higgins, T. J. Gibson, CLUSTAL W: improving the sensitivity of progressive multiple sequence alignment through sequence weighting, position-specific gap penalties and weight matrix choice. Nucleic Acids Res 22, 4673–4680 (1994).

79. X. Robert, P. Gouet, Deciphering key features in protein structures with the new ENDscript server. Nucleic Acids Res 42, W320–324 (2014).

80. E. Beitz, TEXshade: shading and labeling of multiple sequence alignments using LATEX2 epsilon. Bioinformatics 16, 135–139 (2000).

81. Y. Zhang, J. Skolnick, Scoring function for automated assessment of protein structure template quality. Proteins 57, 702–710 (2004).

82. J. Mistry et al., Pfam: The protein families database in 2021. Nucleic Acids Res 49, D412–D419 (2021).

83. S. R. Eddy, Accelerated Profile HMM Searches. PLoS Comput Biol 7, e1002195 (2011).

84. S. Capella-Gutierrez, J. M. Silla-Martinez, T. Gabaldon, trimAl: a tool for automated alignment trimming in large-scale phylogenetic analyses. Bioinformatics 25, 1972–1973 (2009).

85. B. Q. Minh et al., IQ-TREE 2: New Models and Efficient Methods for Phylogenetic Inference in the Genomic Era. Mol Biol Evol 37, 1530–1534 (2020).

86. D. T. Hoang, O. Chernomor, A. von Haeseler, B. Q. Minh, L. S. Vinh, UFBoot2: Improving the Ultrafast Bootstrap Approximation. Mol Biol Evol 35, 518–522 (2018).

87. S. Guindon et al., New algorithms and methods to estimate maximum-likelihood phylogenies: assessing the performance of PhyML 3.0. Syst Biol 59, 307–321 (2010).

88. I. Letunic, P. Bork, Interactive Tree Of Life (iTOL) v5: an online tool for phylogenetic tree display and annotation. Nucleic Acids Res 49, W293–W296 (2021).

