## Supplementary Materials for "Differential higher-order superassembly of HECT-type UBE3 ligases controlled by calcium signals"

**The PDF file includes:**

Figs. S1 to S10
Tables S1

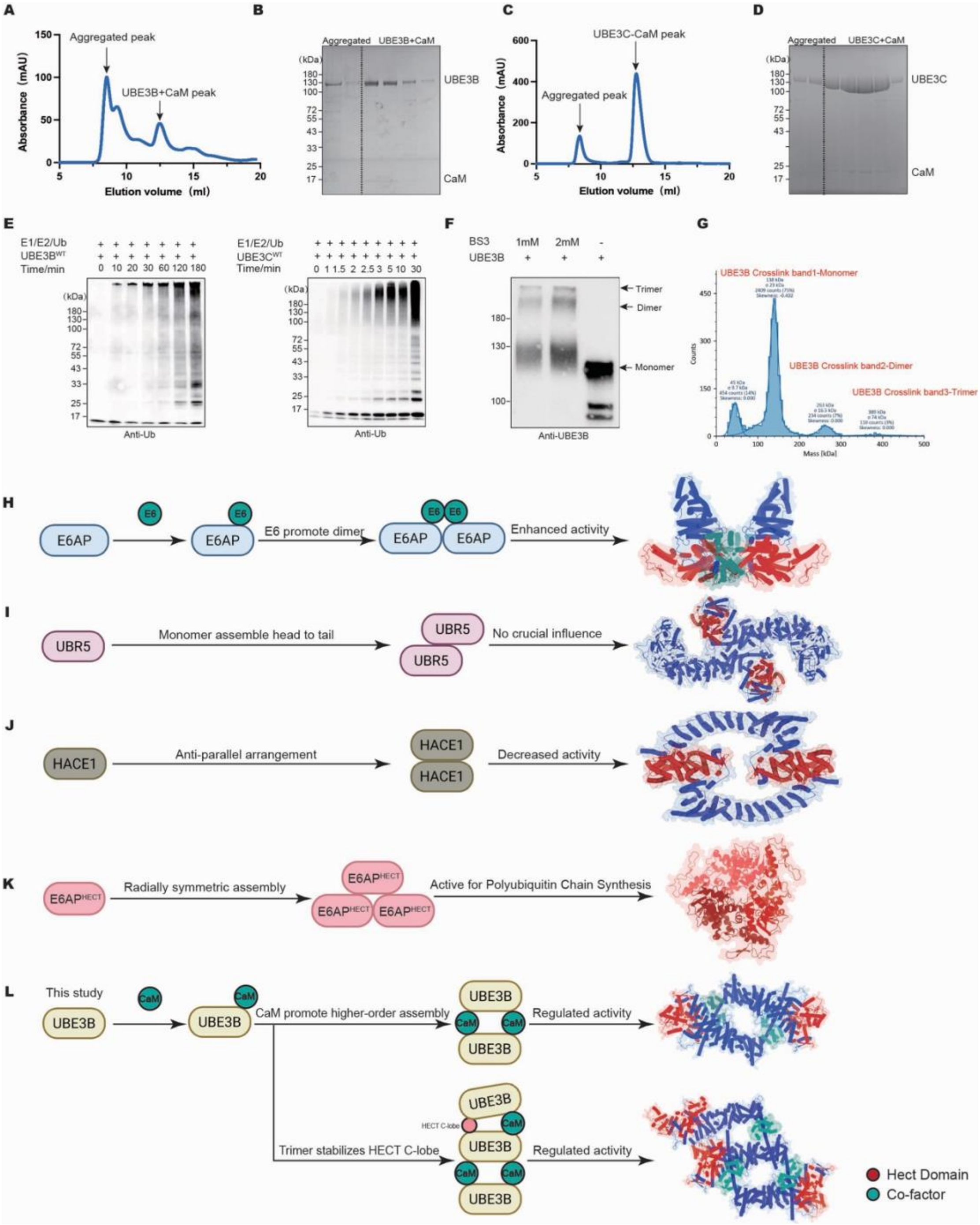


**Fig. S1. Biochemical characterization of UBE3B and UBE3C complexes and comparative analysis of HECT E3 ligase oligomerization.** (**A** and **C**) Size-exclusion chromatography (SEC) profile of the purified UBE3B–CaM (A) and UBE3C–CaM (C) complexes showing separation from aggregated fractions. **(B and D)** Coomassie blue-stained SDS-PAGE analysis confirming the co-purification of UBE3B–CaM (B) and UBE3C–CaM (D). (**E**) In vitro ubiquitination assays of wild-type UBE3B (left) and UBE3C (right). **(F)** BS3 crosslinking analysis of UBE3B detected by immunoblotting. The monomer, dimer and trimer species are observed upon treatment with BS3 (1-2 mM). (**G**) Mass photometry of crosslinked UBE3B revealing monomeric (~125 kDa), dimeric (~250 kDa) and trimeric (~375 kDa) species. (**H** to **L**) Comparison of oligomerization mechanisms and functional implications for HECT E3 ligases. (**H**) E6-

induced E6AP (UBE3A) dimerization (PDB: 8JRN). (**I**) UBR5 forms a head-to-tail dimer assembly (PDB: 8BJA). (**J**) HACE1 forms an anti-parallel dimer (PDB: 8HAE). (**K**) E6AP forms a trimeric assembly. (**L**) CaM-mediated oligomerization of UBE3B. CaM promotes the higher-order assembly of UBE3B.

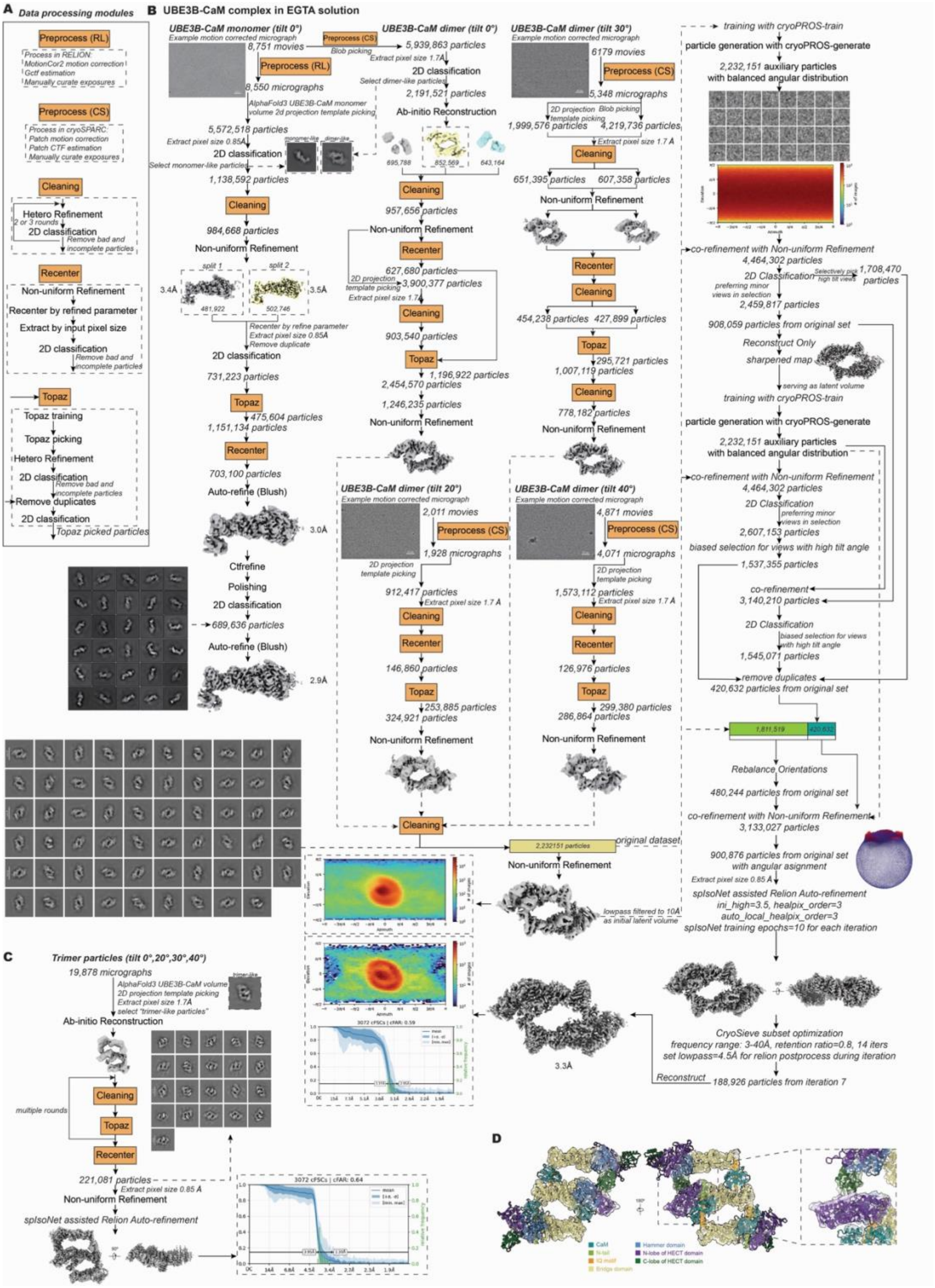


**Fig. S2. Cryo-EM data processing workflow for the UBE3B–CaM complex in the presence of 2 mM EGTA. (A)** Definition of data processing modules. To simplify the workflow, detailed processing steps (Preprocess, Cleaning, Recenter and Topaz) are grouped into functional modules defined here. (**B**) Schematic overview of the cryo-EM processing pipeline, utilizing the modules defined in (A). The workflow spans from representative raw micrographs (collected at 0°, 20°, 30°, and 40° tilt angles) to the final density maps of the UBE3B–CaM monomer and dimer obtained in the presence of 2 mM EGTA. Processing was performed using cryoSPARC and

RELION. Representative intermediate results (2D class averages, 3D classification subsets, directional FSC curves, angular distribution plots) and final reconstructions are shown. (**C**) Identification and data processing workflow for a particle subset representing a putative trimeric assembly. The directional FSC analysis reveals severe preferred orientation and resolution anisotropy. (**D**) Rigid-body fitting of atomic models into the low-resolution map of the putative trimer (shown as a transparent gray surface). A close-up view (right) highlights the T-shaped arrangement of the HECT domain and the fit of the HECT C-lobe in the central UBE3B–CaM protomer.

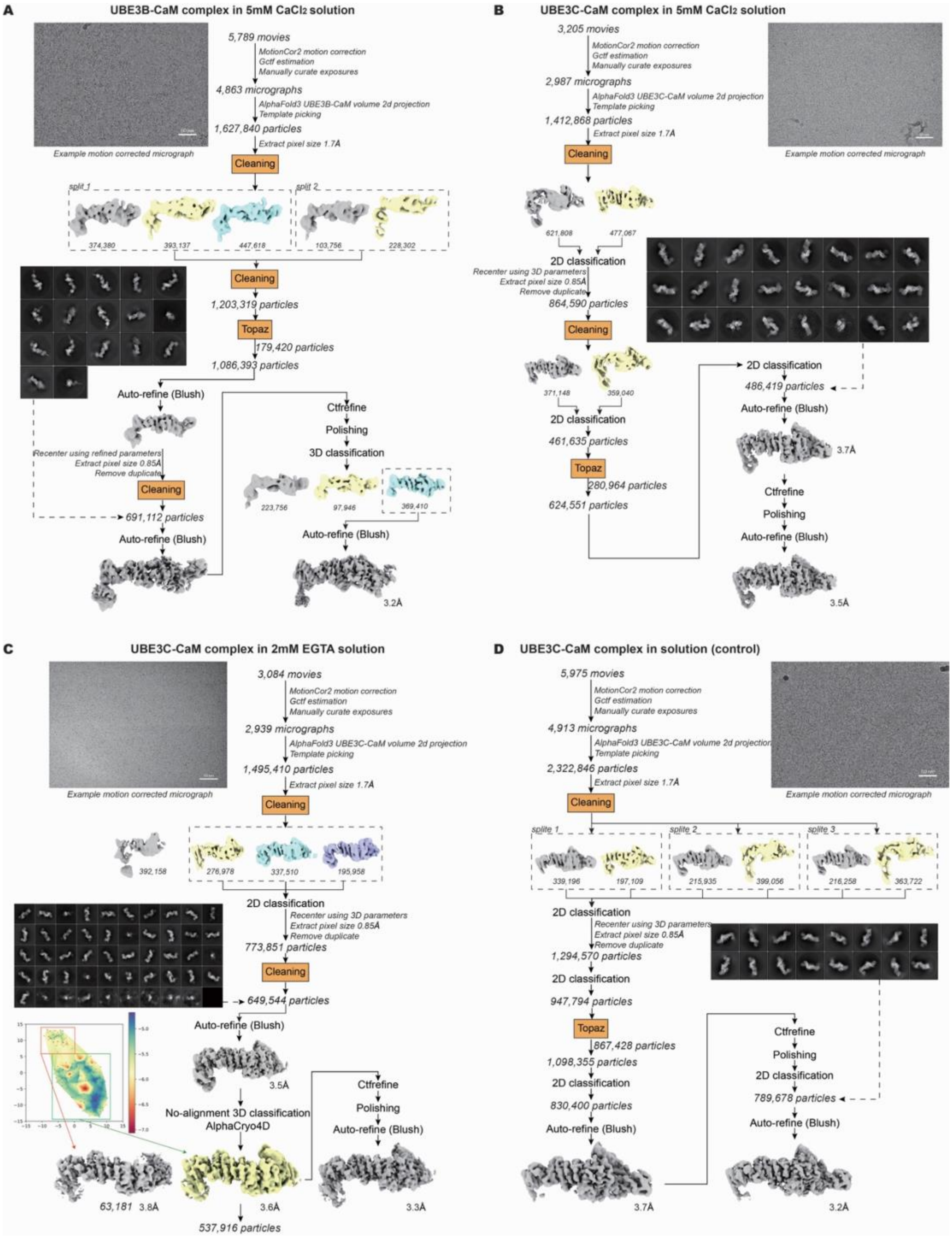


**Fig. S3. Cryo-EM data processing workflows for UBE3B in 5 mM $CaCl_2$ and UBE3C complexes under different calcium conditions.** (**A** to **D**) Schematic overview of the cryo-EM data processing pipelines for the UBE3B–CaM (A) and UBE3C–CaM (B) complexes in the presence of 5 mM $CaCl_2$, as well as the UBE3C–CaM complex in the presence of 2 mM EGTA (C) or in the absence of added $CaCl_2$ or EGTA (control condition) (D). All workflows span from representative raw micrographs to the final density maps, utilizing the data processing modules defined in Extended Data Fig. 2a. For all datasets, particle picking and Topaz training were performed in cryoSPARC, while subsequent classification and refinement steps were carried out in RELION. Representative intermediate results (2D class averages, 3D classification subsets) and final reconstructions are shown.

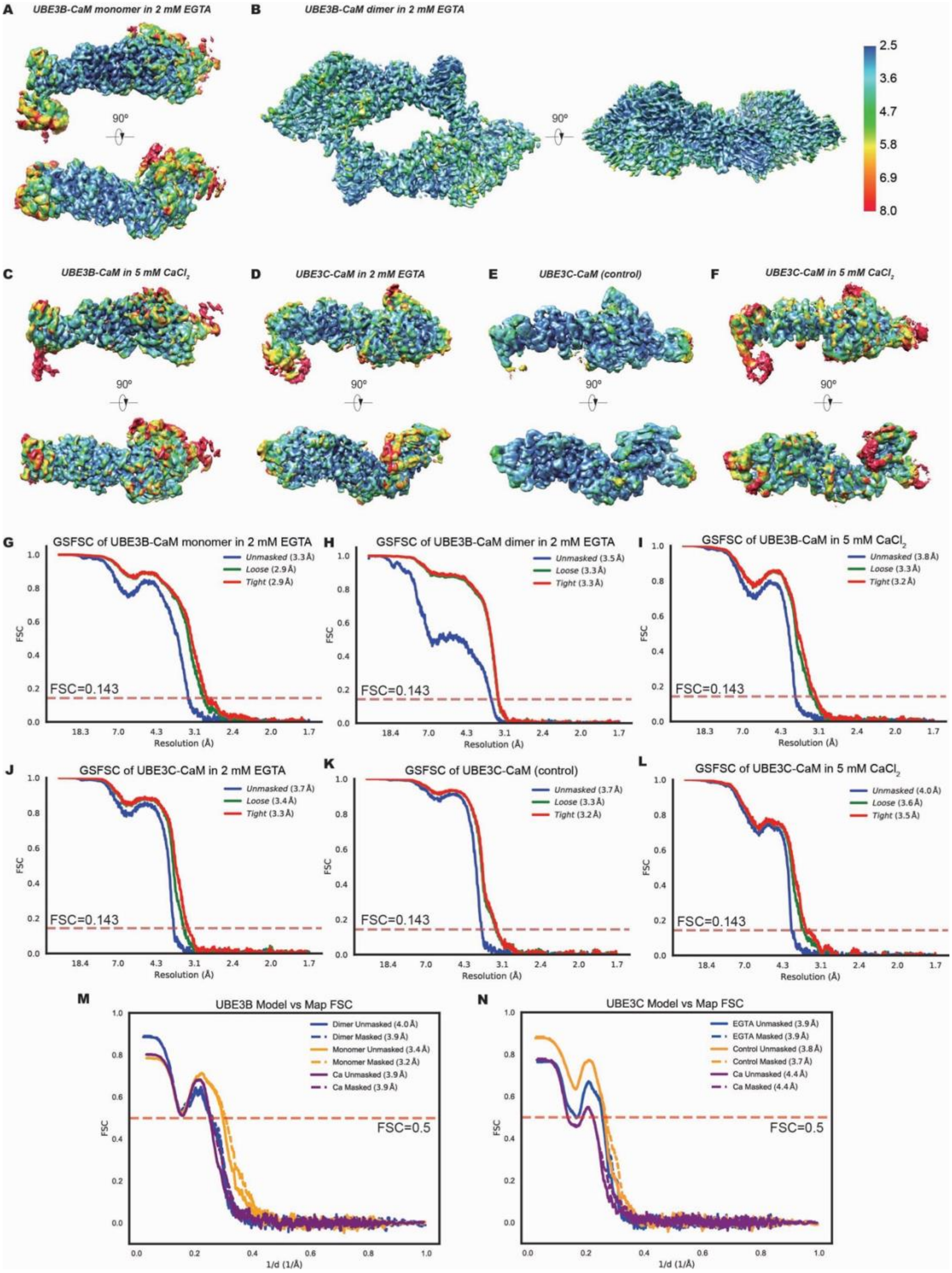


**Fig. S4. Assessment of cryo-EM map and model quality.** (**A** to **F**) Local resolution estimations. The final postprocessed maps are colored according to local resolution estimated by ResMap for the UBE3B–CaM monomer (**A**) and dimer (**B**) in 2 mM EGTA, UBE3B–CaM in 5 mM $CaCl_2$ (**C**) UBE3C–CaM in 2 mM EGTA (**D**) UBE3C–CaM in the control condition (**E**) and UBE3C–CaM in 5 mM $CaCl_2$ (**F**). (**G** to **L**) Gold-standard Fourier shell correlation (FSC) curves for the final reconstructions of the UBE3B-CaM monomer (**G**) and dimer (**H**) in 2 mM EGTA, UBE3B-CaM in 5 mM $CaCl_2$ (**i**), UBE3C-CaM in 2 mM EGTA (**J**), UBE3C-CaM in the control condition (**K**). and UBE3C-CaM in 5 mM $CaCl_2$ (**I**). The FSC curves were calculated using SPIDER, and the reported resolutions are based on the FSC = 0.143 criterion. (**M** and **N**) Map-to-model FSC curves calculated in PHENIX for the UBE3B complexes (**M**) and UBE3C complexes (**N**). Curves correspond to the monomer (EGTA), dimer (EGTA), and $CaCl_2$ datasets for UBE3B, and the EGTA, control, and $CaCl_2$ datasets for UBE3C, as indicated in the legends. The resolution at FSC = 0.5 is indicated.

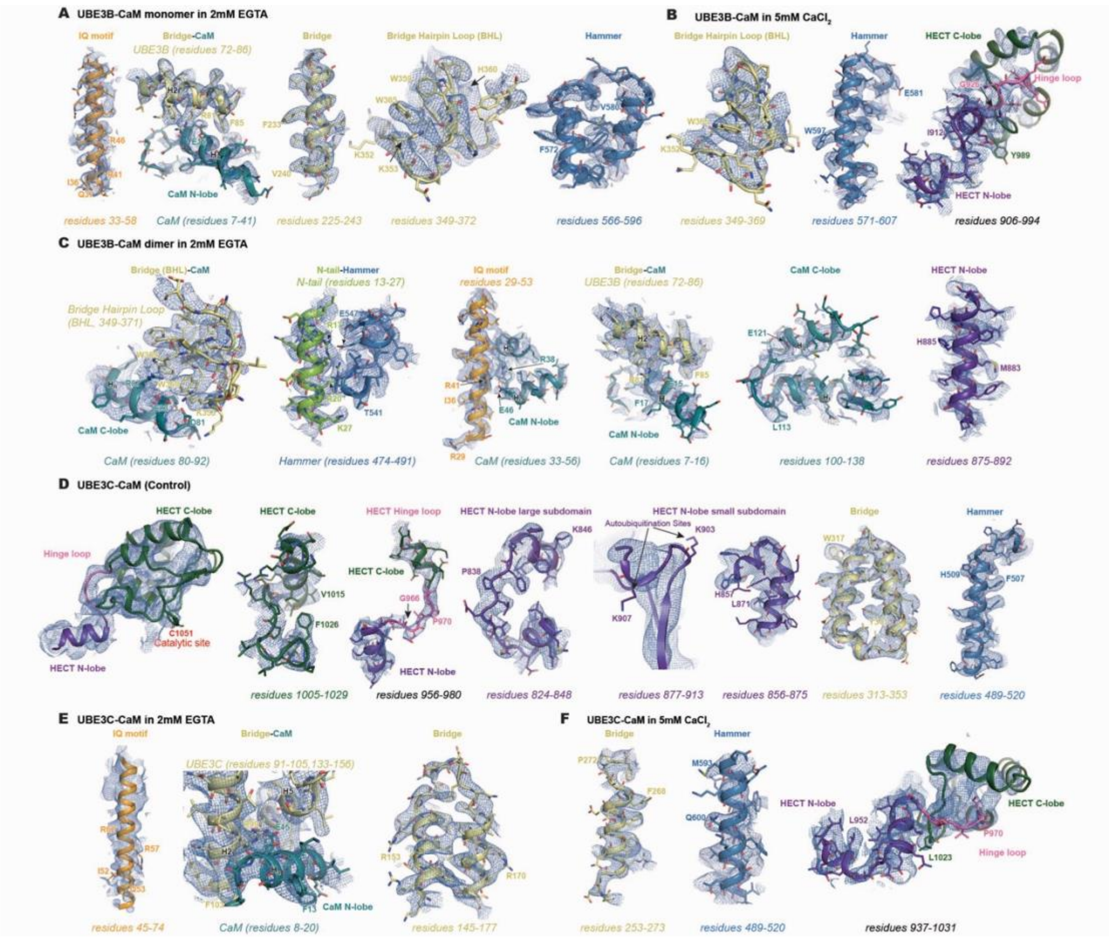


**Fig. S5. Cryo-EM map quality and fitting of key structural elements.**
Representative cryo-EM densities (mesh) and atomic models (sticks) are shown (e.g., IQ motif, Bridge domain, HECT domain, CaM) and specific residue ranges are labeled to highlight the quality of side-chain densities. (**A** to **C**) Representative views of UBE3B–CaM complexes. (A) Monomer (2 mM EGTA) showing IQ motif, BHL helix, Bridge domain, and the intramolecular interface between the Bridge and apo-CaM N-lobe. (B) Monomer (5 mM $CaCl_2$) showing BHL, Hammer, and HECT N- and C-lobes. (C) Dimer (2 mM EGTA) showing BHL–CaM C-lobe interface, the N-terminal tail insertion into the Hammer pocket of the adjacent protomer, the IQ–apo-CaM N-lobe, the intramolecular Bridge–apo-CaM N-lobe interface, and the HECT N-lobe. (**D** to **F**) Representative views of UBE3C–CaM complexes. (D) The complex in the control condition showing the HECT hinge loop, lobes, Bridge, and Hammer domains. (E) The complex in 2 mM EGTA showing the Bridge domain and its interaction with the apo-CaM N-lobe. (F) The complex in 5 mM $CaCl_2$. Shown are the Bridge, Hammer, and HECT C-lobe.

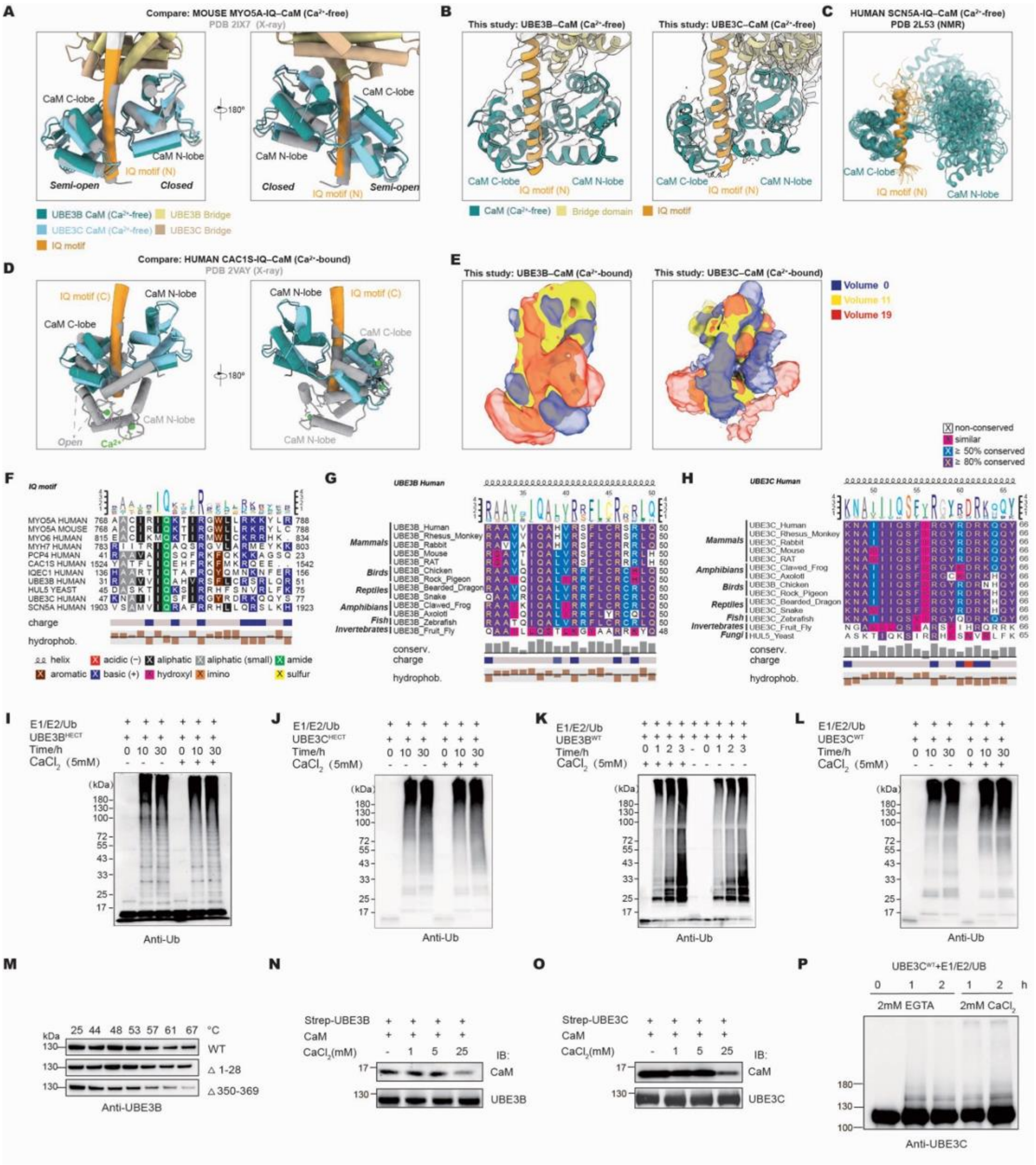


**Fig. S6. Structural comparison of CaM binding and $Ca^{2+}$-dependent regulation.** All structural models are aligned on the IQ motif. **(A)** Comparison of $Ca^{2+}$-free mouse CaM–MYO5A (PDB 2IX7) with $Ca^{2+}$-free UBE3B–CaM and UBE3C–CaM. (**B**) Models of UBE3B and UBE3C in complex with $Ca^{2+}$-free CaM fitted into cryo-EM density maps. **(C)** NMR structures of human SCN5A IQ–CaM ($Ca^{2+}$-free) in solution (PDB 2L53). (**D**) Comparison of the human $Ca^{2+}$-bound CaM–CAC1S (PDB 2VAY) with $Ca^{2+}$-free UBE3B–CaM and UBE3C–CaM. (**E**) 3DVA of $Ca^{2+}$-bound CaM–IQ density in UBE3B (left) and UBE3C (right) monomers (5 mM $CaCl_2$). Representative volumes (#0, #11, #19) are superimposed (blue/yellow/red). (**F** to **H**) Multiple sequence alignment (MSA) of IQ motifs. (F**)** Human UBE3B/UBE3C and other CaM-binding proteins, colored according to residuce properties. (G and H) UBE3B (G) and UBE3C (H) orthologs colored by conservation. (**I** and **J**) Ubiquitin assays comparing FL vs HECT domains of UBE3B (I) and UBE3C (J) under indicated $Ca^{2+}$ conditions.

(**K** and **L**) Time-course ubiquitination assays of $UBE3B^{WT}$ (K) and $UBE3C^{WT}$ (L) (5 mM $CaCl_2$ vs 2 mM EGTA). (**M**) **Cellular thermal shift assays of $UBE3B^{WT}$ and truncation mutants.** (**N** and **O**) $Ca^{2+}$-dependent pulldowns of Strep-tagged UBE3B (N) or UBE3C (O). (**P**) *In vitro* autoubiquitination assay of $UBE3C^{WT}$.

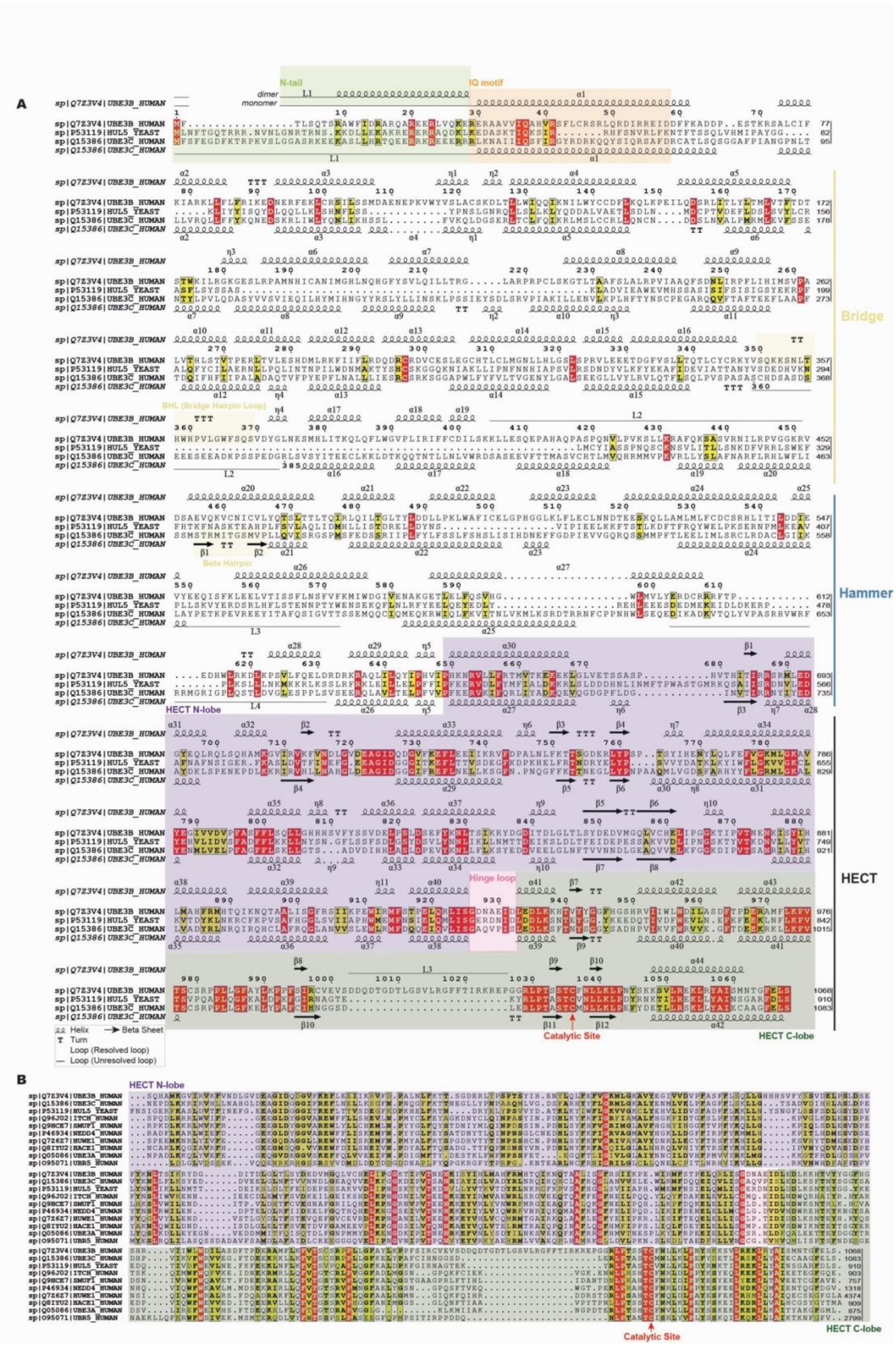

**Fig. S7. Sequence alignment and secondary structure visualization of human UBE3B, UBE3C and yeast HUL5. (A)** Global sequence alignment of full-length human UBE3B (Uniprot Q7Z3V4), human UBE3C (Q15386), and yeast HUL5

(P53119). Secondary structural elements, derived from the cryo-EM structures, are displayed above or under the sequences. Key structural domains (Hammer, Bridge, HECT) are annotated to highlight divergent regions and conserved motifs. (**B**) Multiple sequence alignment of the HECT domains from UBE3B, UBE3C, HUL5, and other representative HECT-family E3 ligases. The catalytic cysteine residues are labeled.

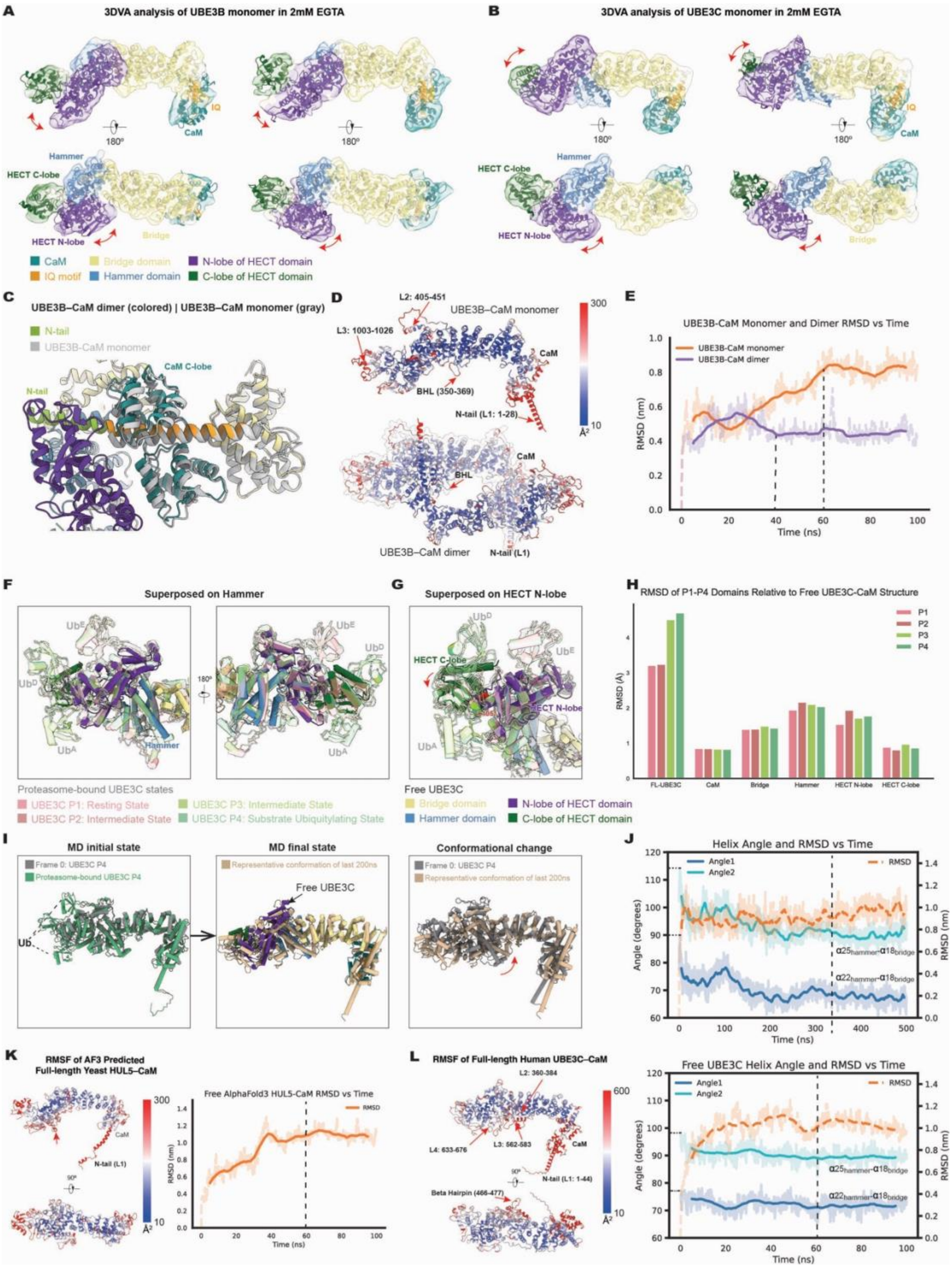


**Fig. S8. Conformational dynamics and structural stability of UBE3B and UBE3C.** (**A** and **B**) 3DVA of UBE3B (A) and UBE3C (B) monomers (2 mM EGTA) showing HECT domain motion and rotation. (**C**) Structural superposition of the UBE3B–CaM dimer (colored) and monomer (gray). (**D** and **E**) MD simulations assessing UBE3B stability. (**D**) Per-residue RMSF mapped onto the UBE3B monomer (top) and dimer (bottom). Colors (blue to red) indicate flexibility. (**E**) RMSD trajectories of UBE3B in monomeric (orange) and dimeric (purple) forms over 100 ns. (**F** to **H**) Comparison of free UBE3C with proteasome-bound states (P1-P4).

**(F)** Superposition aligned on the Hammer domain. P1-P4 states are displayed as transparent cartoons. Ubiquitin are labeled $Ub^A$, $Ub^D$, and $Ub^E$. (**G)** Superposition aligned on the HECT N-lobe. (**H**) RMSD values of P1-P4 states relative to free UBE3C. (**I** and **J**) MD simulation of UBE3C relaxation from P4 state (500 ns). (**I**) Snapshots: initial P4 (left); final frame (gold) superposed on free UBE3C (middle); and comparison of initial vs. relaxed states (right). (**J**) RMSD (orange) and inter-helical angle (blue/teal) trajectories. (**K**) MD of AF3-predicted yeast HUL5–CaM. (**L**) MD of free human UBE3C showing RMSF (left) and trajectories (right).

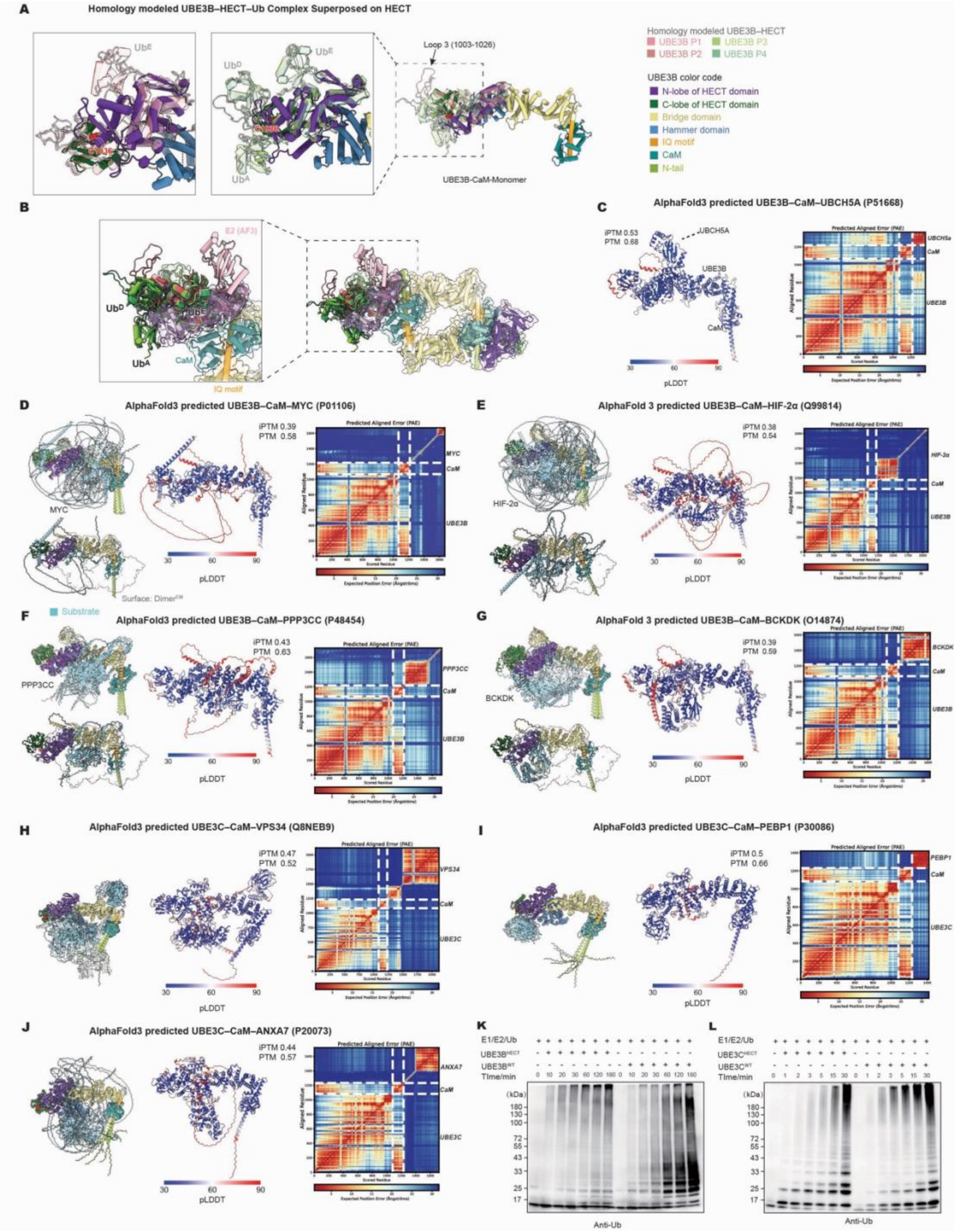


**Fig. S9. Structural insights into UBE3B catalytic architecture and substrate recognition. (A)** Homology modeling of the UBE3B HECT–Ub complexes based on proteasome-bound UBE3C templates ($Ub^{A}$, $Ub^{D}$, and $Ub^{E}$). Models are superposed onto the UBE3B monomer (colored by domain). **(B)** Superposition of the modeled UBE3B HECT–Ub complexes onto the UBE3B dimer and the AF3-predicted UBE3B–CaM–E2 complex (showing E2 only). (**C**) AF3-predicted structure of the UBE3B–CaM–UBCH5A (E2) complex. Model colored by pLDDT (blue: low; red: high); PAE matrix shown on right. (**D** to **G**) AF3-predicted UBE3B–substrate interactions. Superposition of ten independent AF3 predictions (left top) and a representative predicted model superposed onto the UBE3B dimer, which is rendered as a transparent surface (left bottom) with the substrates colored blue, pLDDT, and PAE (right). Substrates: (D) MYC; (E) HIF-2α; (F) PPP3CC; and (G) BCKDK. (**H** to **J**) AF3-predicted UBE3C–substrate interactions (layout as in D–G). Substrates: (H) VPS34; (I) PEBP1; and (J) ANXA7. (**K** and **L**) *In vitro* ubiquitin assays comparing UBE3B$^{WT}$ vs UBE3B$^{HECT}$ (K) and UBE3C$^{WT}$ vs UBE3C$^{HECT}$ (L) (anti-Ub immunoblot).

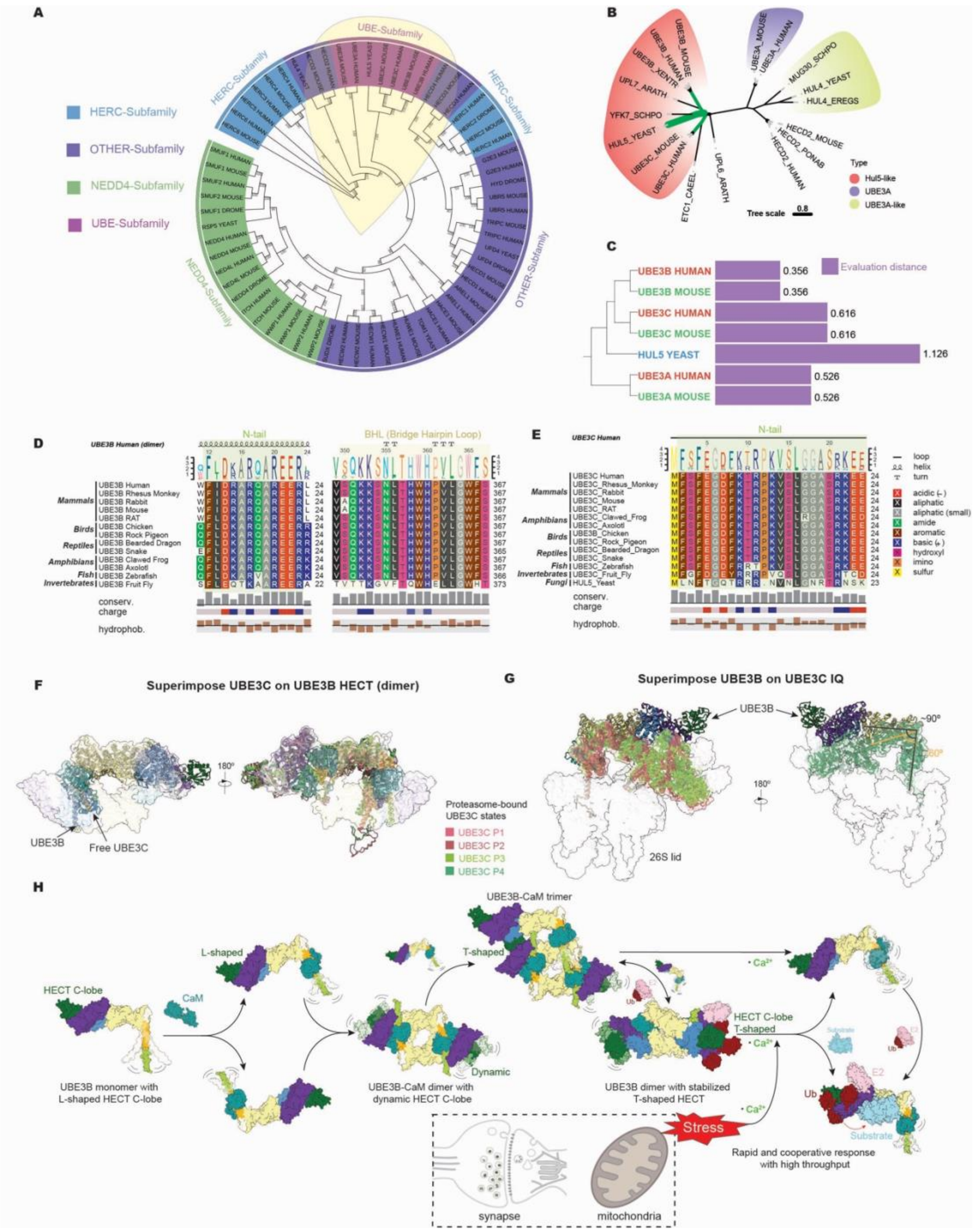


**Fig. S10. Evolutional and structural divergence of UBE3B and UBE3C. (A)** Phylogenetic tree of HECT E3 ubiquitin ligases (human, mouse, fly, yeast). (**B**) Linear phylogenetic tree shows the evolutionary relationships among human and mouse UBE3A, UBE3B, and UBE3C, and yeast HUL5. (**C**) Unrooted phylogenetic tree of UBE3A/UBE3A-like and HUL5-like proteins across diverse species. (**D** and **E**) Sequence alignments of the dimer-associated N-tail (11-24) and BHL motif in UBE3B (D) and proteasome binding associated N-terminal strand (1-24) in UBE3C (E) across diverse species. (**F**) Superposition of UBE3C states onto the UBE3B–CaM dimer (aligned on the HECT domain). (**G**) Superposition of UBE3B–CaM monomer onto the proteasome-bound UBE3C (aligned on the IQ motif). Angles between IQ-helix and Bridge domain are indicated (~90° for UBE3B, black; ~60° for UBE3C, orange). (**H**) Proposed mechanistic model. In the absence of $Ca^{2+}$, CaM stabilizes UBE3B dimers (L-shaped HECT). Trimer formation induces an active T-shaped HECT conformation. Elevated $Ca^{2+}$ triggers dissociation into monomers, exposing substrate binding sites for ubiquitination.

**Table S1. Cryo-EM data collection, refinement and validation statistics**

| | UBE3B dimer (2 mM EGTA) | UBE3B monomer (2 mM EGTA) | UBE3B (5 mM $Ca^{2+}$) | UBE3C (2 mM EGTA) | UBE3C (control) | UBE3C (5 mM $Ca^{2+}$) |
|---|---|---|---|---|---|---|
| **Data collection and processing** | | | | | | |
| Magnification | 105,000 | 105,000 | 105,000 | 105,000 | 105,000 | 105,000 |
| Voltage (kV) | 300 | 300 | 300 | 300 | 300 | 300 |
| Electron exposure ($e^-$/Å$^2$) | 60 | 60 | 60 | 60 | 60 | 60 |
| Defocus range (μm) | -0.5 to -2.5 | -0.5 to -2.5 | -0.5 to -2.5 | -0.5 to -2.5 | -0.5 to -2.5 | -0.5 to -2.5 |
| Pixel size (Å) | 0.425 | 0.425 | 0.425 | 0.425 | 0.425 | 0.425 |
| Symmetry imposed | C1 | C1 | C1 | C1 | C1 | C1 |
| Initial particle images (no.) | 6,662,996 | 1,138,592 | 1,627,840 | 1,495,410 | 2,322,846 | 1,412,868 |
| Final particle images (no.) | 188,926 | 689,636 | 369,410 | 537,916 | 789,678 | 486,419 |
| Map resolution (Å) | 3.3 | 2.9 | 3.2 | 3.3 | 3.2 | 3.5 |
| FSC threshold | 0.143 | 0.143 | 0.143 | 0.143 | 0.143 | 0.143 |
| Map resolution range (Å) | 2.5-8.0 | 2.5-8.0 | 2.5-8.0 | 2.5-8.0 | 2.5-8.0 | 2.5-8.0 |
| **Refinement** | | | | | | |
| Initial model used (AphaFold3-AF3) | AF3 UBE3B-CaM | AF3 UBE3B-CaM | AF3 UBE3B | AF3 UBE3C-CaM | AF3 UBE3C | AF3 UBE3C |
| Model resolution (Å) | 3.9 | 3.2 | 4.0 | 3.9 | 3.7 | 4.4 |
| FSC threshold | 0.5 | 0.5 | 0.5 | 0.5 | 0.5 | 0.5 |
| Model resolution range (Å) | 2.5-6.0 | 2.5-6.0 | 2.5-8.0 | 2.5-8.0 | 2.5-6.0 | 2.5-8.0 |
| Map sharpening *B* factor (Å$^2$) | 0 | -50 | -50 | -50 | -50 | -50 |
| **Model composition** | | | | | | |
| Non-hydrogen atoms | 16242 | 8958 | 7728 | 8673 | 7375 | 7410 |
| Protein residues | 1996 | 1106 | 950 | 1077 | 914 | 918 |
| Ligands | 0 | 0 | 0 | 0 | 0 | 0 |
| *B* factors (Å$^2$) | 43.58 | 109.99 | 205.29 | 169.13 | 179.37 | 120.99 |
| **R.m.s. deviations** | | | | | | |
| Bond lengths (Å) | 0.003 | 0.003 | 0.003 | 0.003 | 0.003 | 0.003 |
| Bond angles (°) | 0.486 | 0.584 | 0.584 | 0.63 | 0.556 | 0.593 |
| **Validation** | | | | | | |
| MolProbity score | 1.57 | 1.46 | 1.65 | 1.68 | 1.37 | 1.6 |
| Clashscore | 8.31 | 5.57 | 7.65 | 8.78 | 4.53 | 6.33 |
| Rotamers outliers (%) | 0.49 | 0.7 | 0.46 | 0.83 | 0.24 | 0.85 |
| **Ramachandran plot** | | | | | | |
| Favored (%) | 97.42 | 97.09 | 96.5 | 96.71 | 97.23 | 96.24 |
| Allowed (%) | 2.48 | 2.82 | 3.5 | 3.29 | 2.65 | 3.76 |
| Outliers (%) | 0.10 | 0.09 | 0 | 0 | 0.11 | 0 |
